# Alternatively activated monocyte-derived myeloid cells promote extracellular pathogen persistence within pulmonary fungal granulomas

**DOI:** 10.1101/2025.05.23.655817

**Authors:** Yufan Zheng, Makheni Jean Pierre, Eduard Ansaldo, Hannah E. Dobson, Olena Kamenyeva, Chinaemerem U. Onyishi, Eric V. Dang

**Affiliations:** Molecular Mycology and Immunity Section, Laboratory of Host Immunity and Microbiome, National Institute of Allergy and Infectious Diseases, National Institutes of Health, Bethesda, MD, 20892, USA; Department of Microbiology and Immunology, Georgetown University, Washington, DC, 20057, USA; Immunology Graduate Group, Perelman School of Medicine, University of Pennsylvania, Philadelphia, PA, 19104, USA; Research Technology Branch, Biological Imaging Section, National Institute of Allergy and Infectious Diseases, National Institutes of Health, Bethesda, MD, 20892, USA

**Author notes:** Correspondence (E.V.D).

## Abstract

Inhaled fungal pathogens often generate granuloma-contained latent infections that can reactivate to cause invasive disease. However, the mechanisms underlying the inability to generate sterilizing immunity against latent infection remains poorly understood. Here, we leveraged spatial transcriptomics and flow cytometry to characterize the immune dynamics and cellular architecture of cryptococcal granulomas. Using fate mapping and murine genetic tools, we demonstrate that alternative activation of monocyte-derived myeloid cells by CD4^+^ T helper 2 cells antagonizes pulmonary fungal clearance during latent infection. In contrast to the prevailing view in the field, we find that alternatively activated myeloid cells are not an intrinsic replication niche for the fungus and more broadly, *Cryptococcus* predominantly resides in the extracellular environment. We propose a T helper 2 cell-myeloid circuit establishes a local immunosuppressive environment to drive extracellular fungal persistence, which could be leveraged as a new target for host-directed therapy to treat latent fungal infections.

## Main

A key feature of many fungal pathogens that cause human disease is their ability to drive latent, persistent infections^1^. Fungi such as *Cryptococcus*, *Histoplasma*, and *Coccidioides*, can establish latent infections contained within pulmonary granulomas^2–4^, micro-anatomical structures composed of heterogenous immune cells^5^. Although asymptomatic in healthy individuals, these infections can reactivate to cause invasive, disseminated disease in patients with suppressed immune systems^6^. For instance, *Cryptococcus neoformans* causes asymptomatic latent pulmonary infections in immunocompetent people but life-threatening meningitis in patients with HIV infection, which accounts for 19% of AIDS-related mortality^7^.

Human patients with auto-antibody production against interferon gamma (IFNγ) are more susceptible to *Cryptococcus* infection, and mice genetically deficient in IFNγ or treated with IFNγ blocking antibodies succumb faster to serotype D infections^8–11^, suggesting IFNγ responses are induced in human and murine *Cryptococcus* infection and required for protection. While it is clear that IFNγ is capable of driving sterilizing immunity in certain contexts, as *C. neoformans* strains that overexpress IFNγ are rapidly cleared from mice and IFNγ is required for host protection in vaccine responses^12,13^, why this response cannot drive sterilizing immunity during chronic infection remains unknown. Whether this is due to inefficient protective responses (i.e. exhaustion of IFNγ-producing lymphocytes) or active immunosuppressive circuits is a pressing question.

In acute murine infections with *Cryptococcus* serotype A and D, which are uniformly lethal in immunocompetent C57BL/6J (B6) mice, the immune response is overwhelmingly biased towards type 2 inflammation, characterized by T helper (T_H_) 2 cells, eosinophilia, and alternative activation of macrophages^14–16^. Deletion of type 2 signaling in this setting results in decreased fungal burden in the lungs^17,18^. On the other hand, SJL/J mice are resistant to acute *Cryptococcus* infection, thus better approximating human infection, and show a dominant type 1 response with little induction of type 2 cytokines^9^. It is, therefore, unclear whether the induction of type 2 inflammation in *Cryptococcus* infection is restricted to settings of murine acute lethal infections that do not appropriately mimic human disease progression.

Granulomas are microanatomical structures that can restrict microbial dissemination, but also generate an environment that allows pathogen persistence^5^. In zebrafish *Mycobacterium marinum* infection, recruitment of macrophages into the granuloma supports bacterial persistence by providing a continually renewed bacterial replication niche^19–22^. Additionally, in zebrafish, STAT6 signaling induces macrophage epithelioid transformation via upregulation of E-cadherin to form a myeloid ring that functions to prevent dissemination^23,24^. In mice, however, myeloid cell-specific deletion of IL-4Rα has no phenotype with *Mycobacterium tuberculosis* (Mtb) infection^25^, whereas in the macaque mycobacterial infection model, type 2 immunity has been associated with higher bacterial burdens^26^. While the role of type 2 immunity in the Mtb granuloma is still controversial, other potential barriers to sterilizing immunity in this context could be limited antigen presentation, TGF-β signaling, and T cell exhaustion^27–29^. The precise definition of a granuloma remains elusive though at a minimum, there must be spatial organization of the cells that comprise the structure^5^. In Mtb granulomas, there is an inner core comprised mainly of macrophages and granulocytes that is surrounded by a lymphocyte cuff, consistent with spatially distinct chemokine expression patterns found in transcriptomics studies. While fungal granulomas are commonly found in human lung biopsies^30^, we know very little about the organization of these structures. Dissecting how immune responses and involved immunocytes control protective versus immunosuppressive outcomes within granulomas is important, as this will reveal the immunological circuits that antagonize sterilizing immunity. Mechanistic understanding has traditionally been limited by a dearth of animal models that generate fungal granulomas. However, it is reported that C57BL6/J mice infected with a *C. neoformans* strain lacking glucosylceramide synthase (Δ*gcs1*) display robust granuloma formation^31^. Murine infection with Δ*gcs1* results in a chronic, asymptomatic infection that can reactivate upon blockade of lymphocyte recirculation^32^. This opens the door to utilizing B6 mouse genetics to probe protective versus detrimental immunity in a pulmonary granuloma model with a genetically tractable pathogen^2,33^.

In this study, we leveraged the Δ*gcs1* granuloma model to uncover the mechanisms underlying fungal persistence during chronic infection. Using spatial transcriptomics, we observed that fungal granulomas show an architecture similar to human and non-human primate Mtb granulomas with an inner myeloid core surrounded by a T/B-lymphocyte cuff. Using flow cytometry, we found that the immune system has an early peak of type 2 inflammation that wanes but persists throughout the infection course. While wild-type mice generally did not succumb to infection, they maintained persistent pulmonary load out to at least 5 months, which was lost in *Il4/Il13*^-/-^ mice. T_H_2 cells appear to be the main producers of type 2 cytokine during granulomatous infection, as ablation of GATA3 from post-thymic lymphocytes resulted in pulmonary fungal clearance, whereas deletion of innate lymphocytes 2 (ILC2s), eosinophils, or basophils had no such effects. Production of IL-4 and IL-13 led to the alternative activation of myeloid (AAMs) cells exclusively derived from recruited monocytes, as evidenced by *Ms4a3*-fate mapping. Deleting STAT6 from monocyte-derived cells reduced pulmonary fungal load. While many studies have suggested that AAMs are detrimental during infection by providing an intracellular replication niche, we found that *Cryptococcus* exists almost entirely in the extracellular space and STAT6-mixed chimeras showed no intrinsic differences in myeloid fungal load. We further found that STAT6-deficient macrophages were able to localize closer to *Cryptococcus* yeasts, indicating a positioning effect by type 2 cytokines. Collectively, our findings suggest that type 2 cytokines actively suppress the capacity of myeloid cells to mediate extracellular fungal clearance in pulmonary granulomas.

## Results

### B6 mice can control KN99*Δgcs1* to form a latent infection contained by pulmonary granulomas

To investigate the mechanisms underlying *Cryptococcus* latency, we sought to establish latent infection in C57BL/6 mice, the background on which most murine genetic tools exist. Consistent with previous reports, wild type (KN99α) *Cryptococcus neoformans* induced a lethal infection, where we could not observe clear protective roles for IFNγ (*Ifng^-/-^*) or adaptive immunity (*Rag1^-/-^*) (Extended Data Fig. 1a)^9^. Moreover, even *Il4/Il13^-/-^*mice had a mortality rate as high as 70% (Extended Data Fig. a). Rather than displaying anatomical confinement, wild type KN99α *Cryptococcus* infection was diffuse across the pulmonary tissue (Extended Data Fig. 1b). Additionally, we found that IFNγ production by lymphocytes was unable to be maintained as infection progressed, and iNOS production by myeloid cells could not be clearly observed at any timepoint during lethal infection (Extended Data Fig. 1c and d), demonstrating a failure to induce protection in wild type hosts, which contrasts with human *Cryptococcus* infection where immunocompetent hosts normally do not succumb to infection.

On the other hand, Δ*gcs1 Cryptococcus* was reported to be capable of inducing a latent infection in B6 mice^31^. Using Δ*gcs1* on a KN99α background, we found that murine infection with this strain resulted in organized granulomas in B6 mouse lungs starting at 35 days post infection (dpi) (Fig. 1a). We performed thick-tissue imaging with CD11c^Yfp^ reporter mice infected with a Δ*gcs1* mCherry fluorescent reporter strain to visualize localization of myeloid cells and yeast during latent infection. We observed CD11c^+^ cells surrounding *C. neoformans* and forming a spatially confined structure (Fig. 1b).

**Fig. 1:**
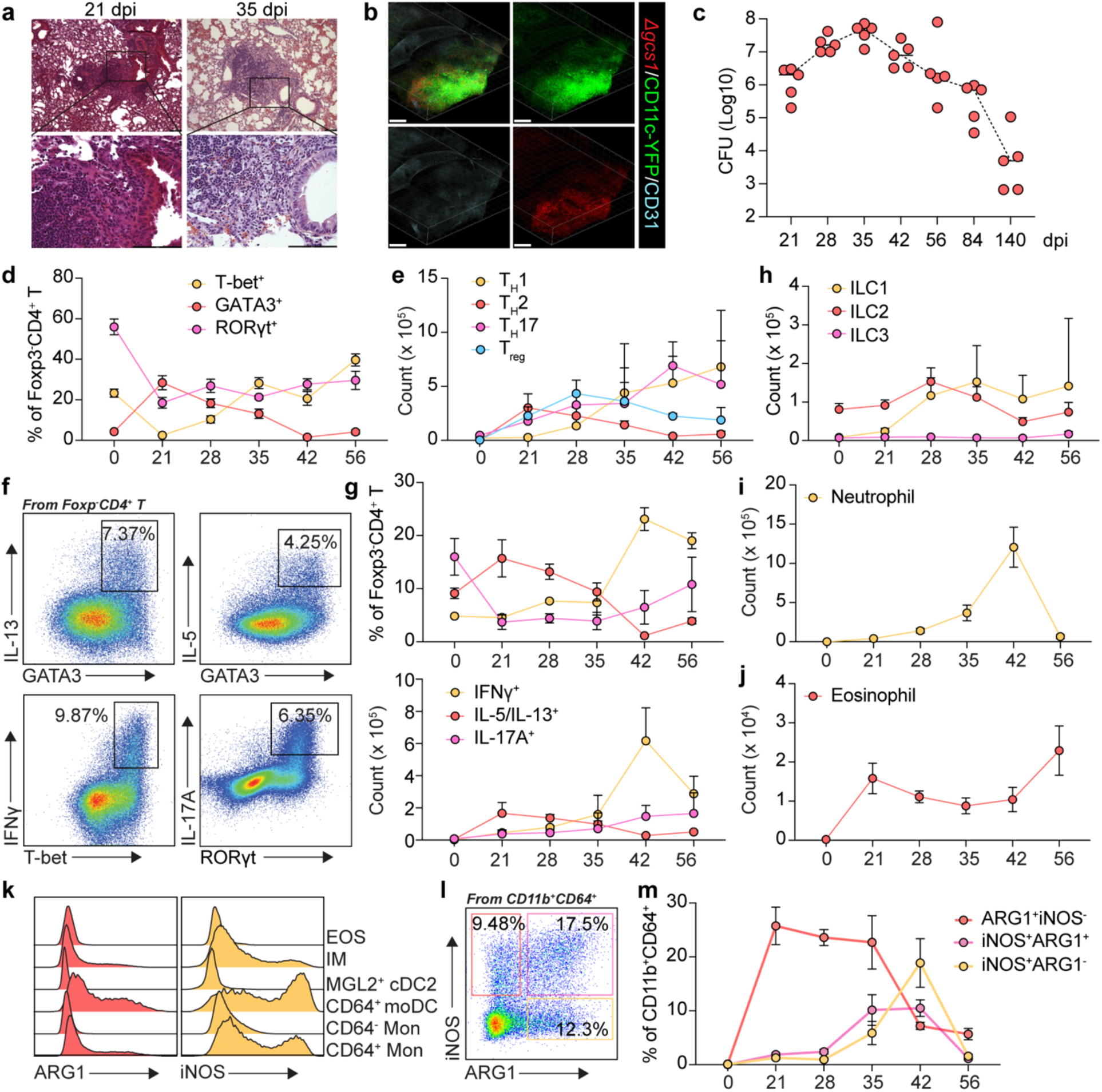
Distinct waves of type 1 and type 2 immune responses during cryptococcus latent infection. (a) Representative images for H&E staining of lung tissues from mice infected with KN99 *Δgcs1* at 21 dpi and 35 dpi. Scale bar, 100 μm. (b) Representative images for thick-tissue imaging of lungs from KN99 *Δgcs1*-infected *CD11c^Yfp^* reporter mice at 35 dpi. Scale bar, 400 μm. (c) Colony-forming unit (CFU) analysis during KN99 *Δgcs1*-infection. (d-m) The dynamics of changes in immune cells during KN99 *Δgcs1* infection (n=5). (d) Statistics for the percent expression of transcription factors, T-bet, GATA3, and RORγt within the Foxp3-CD4+ T cell compartment. (e) Statistics for counts of CD4 subsets. (f) Flow plots for cytokine (IL-13, IL-5, IFNγ, and IL-17A) production in restimulated CD4 T cells. (g) Statistics for the ratio and counts of cytokine-producing CD4 T cells. Statistics for the count of (h) innate lymphocytes, (i) neutrophils, and (j) eosinophils. (k) Histogram of ARG1 and iNOS expression in different types of myeloid cells. (l) Flow plot for ARG1 and iNOS expression in CD64+CD11b+ myeloid cells. (m) Statistics for the ratio of ARG1 and iNOS expression in CD64+CD11b+ myeloid cells. Data are present by mean ± SD. *P < 0.05 and ****P < 0.0001 by unpaired two sided student’s t-test.

To confirm that Δ*gcs1* can drive a chronic infection, we infected B6 mice and collected lungs for colony-forming unit assays (CFUs) across a time course. Pulmonary fungal loads peaked at 35 dpi at which point they began to contract, but CFUs remained detectable out to 140 days (Fig. 1c). To further test if Δ*gcs1* infection was a good mimic of latent infection controlled by adaptive immunity, we administered anti-CD4 antibodies to deplete CD4^+^ T helper cells starting from the peak of infection, 35 dpi (Extended Data Fig. 1e). Using intravascular labeling with anti-CD45-APC, flow cytometry analysis at 49 dpi showed a successful CD4 depletion both in circulation and in the lungs, although there were more residual CD4^+^ T cells in the pulmonary interstitium (Extended Data Fig. 1f and g). Consistent with a protective role for helper T cells, the pulmonary fungal burden in CD4-depleted mice was markedly higher than that in isotype control-treated mice (Extended Data Fig. 1h). Taken together, and consistent with previous reports using this model^31,32^, these data show that Δ*gcs1* induces a latent infection in B6 mice, thus enabling use of genetic mouse tools for further mechanistic studies.

### Distinct waves of type 1 and 2 immune responses characterize *Cryptococcus* latent infection

To investigate immune response dynamics during latent *Cryptococcus* infection, we performed flow cytometry analysis to systematically investigate T cell responses during Δ*gcs1* latent infection (Extended Data Fig. 1i). Consistently, GATA3^hi^ T_H_2 cells peaked in the lungs at 21 dpi, whereas T-bet^+^ T_H_1 cells expanded strikingly from 35 to 42 dpi (Fig. 1 d and e). To assess cytokine production competency, we stimulated T cells with phorbol myristate acetate (PMA) and ionomycin and found that the dynamics of cytokine production generally followed the same kinetics as T cell transcription factor expression (Fig. 1f and g). Intriguingly, we observed strikingly high IFNγ production by Th1 cells at 42 dpi (Fig. 1g), when the pulmonary fungal burdens began to contract (Fig. 1c). We also looked at the dynamics of innate lymphocytes, neutrophils and eosinophils (Fig. 1h-j), which showed the same pattern with an early type 2 wave dominated by eosinophils and a later type 1 wave.

Macrophages and monocytes (Mac/Mon) are innate immune cells that can shape their effector outputs in response to lymphocyte-derived cytokines^34^. While an oversimplification *in vivo*, myeloid cells can be classically activated by type 1 cytokine signaling through STAT1 or alternatively activated by type 2 signaling through STAT6^35^. It has been argued *in vitro* that these two polarization states are mutually antagonistic, whereby STAT6 activation will repress STAT1 target genes (i.e. *Nos2*) and STAT1 will repress STAT6 target genes (i.e. *Arg1*).^36–39^. We performed flow cytometry analysis to assess iNOS and ARG1 expression by different myeloid cells (Extended Data Fig. 2a-c) and found that iNOS and ARG1 were expressed by the same compartments of myeloid cells (CD11b^+^CD64^+^) including IMs, moDCs, and CD64^+^ monocytes (Fig. 1k). However, rather than displaying a mutually exclusive expression pattern, ARG1 was expressed equally by iNOS^+^ and iNOS^-^ cells (Fig. 1l and m). The numbers of type 1 and 2 responsive myeloid cells tracked consistently with T_H_1 and T_H_2 cells; the iNOS expression peaked at 42 dpi (Fig. 1m and Extended Data Fig. 2d), the same as the IFNγ production.

These data strongly argue that induction of type 2 inflammation is not restricted to acute, lethal infection. During chronic infection, type 2 immunity peaked at 21 dpi and then contracted as a type 1 response took over; however, throughout all timepoints there is ongoing type 2 cytokine production and myeloid cell responsiveness.

### Type 2 immunity is detrimental during chronic infection

Taking advantage of this latent infection model in B6 mice, we infected a panel of cytokine-deficient mice to probe the contributions of type 1, 2, and 17 responses to the latent infection (Fig. 2a). Consistent with published data, most WT B6 mice (85%) survived until 140 dpi^31^. IFNγ- and *Rag1*-deficient mice both showed a 100% mortality rate by 64- and 93-days post infection, respectively, demonstrating that type 1 immunity and adaptive lymphocytes in general are required for protection. In contrast, all *Il4/Il13^-/-^*mice survived the infection and had undetectable pulmonary fungal burdens at 140 dpi (Fig. 2b), suggesting that type 2 cytokines antagonize sterilizing immunity during latent infection. To assess the relative contributions of IL-4 vs IL-13, we generated bone marrow chimeras with WT, *Il4^-/-^*, *Il13^-/-^*, and *Il4/Il13^-/-^* donors transplanted into congenic irradiated recipients. After reconstitution and infection, we found that while both single cytokine knockouts resulted in significantly lower fungal burdens at 35 dpi, *Il4^-/-^* had a more striking phenotype, albeit not as strong as the double cytokine knockout, arguing that there is a dominant immunosuppressive role for IL-4 (Fig. 2b and c). ARG1 expression in CD64^+^CD11b^+^ myeloid cells correlated with the fungal burden decrease (Fig. 2d). These data suggest that hematopoietic IL-4 and IL-13 both contribute to fungal persistence to slightly different degrees; this is in line with previous reports that IL-13 receptor expression is more restricted to non-immune cells that would be less likely to influence pathogen burdens compared to the hematopoietic compartment^40^.

**Fig. 2:**
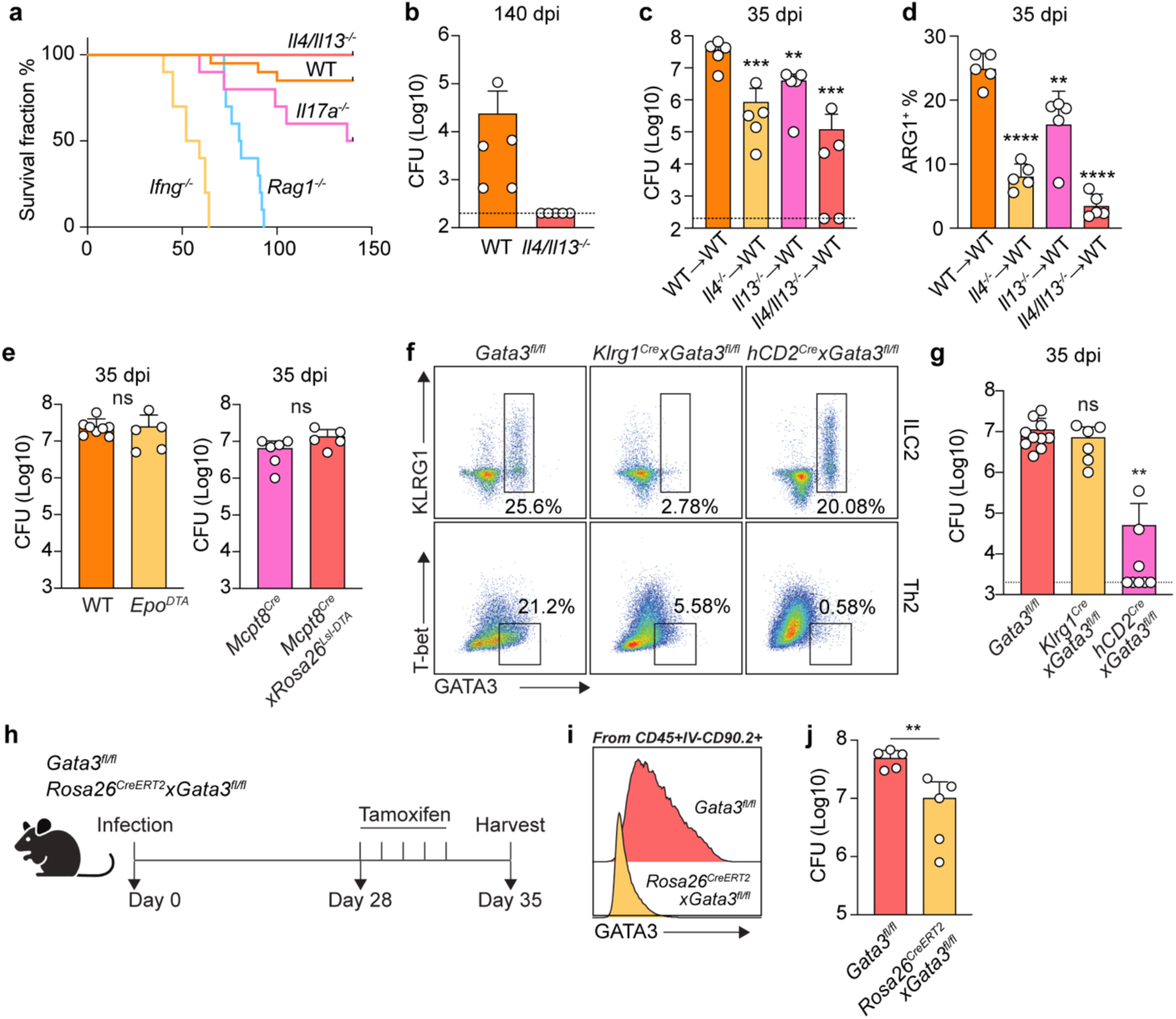
Type 2 immunity is detrimental during *Cryptococcus* chronic infection. (a) Survival rate of mice of the indicated genotypes infected with KN99 *Δgcs1* (n=10). (b) CFU analysis of mice infected with KN99 *Δgcs1*at 140 dpi. (c, d) Bone marrow chimeras from WT, *Il4^-/-^*, *Il13^-/-^*, or *Il4/Il13^-/-^* donor mice. (c) CFU analysis at 35 dpi. (d) ARG1 expression by CD64+CD11b+ myeloid cells at 35 dpi. (e) CFU analysis in eosinophil deficient (Epo^DTA^) or basophil deficient (*Mcpt8^Cre^* x *Rosa26^LSL-DTA^*) mice as well as their littermate controls at 35 dpi. (f) Flow plots for ILC2 and Th2 cells from *Gata3^fl/fl^*, *Klrg1^Cre^ x Gata3 ^fl/fl^*, and *hCD2^Cre^ x Gata3^fl/fl^* mice infected with KN99 at 35 dpi. Plots on the top were gated on CD90.2+TCRβ- and plots on the bottom were gated on CD90.2+TCRβ+Foxp3-CD4+. (g) CFU analysis for the mice in (F) at 35 dpi. (h-j) GATA3 temporal depletion in mice infected with KN99 *Δgcs1*. (h) Scheme for GATA3 temporal depletion experiment. (i) Histogram for GATA3 expression with in CD90+ compartment. (j) CFU analysis. Data are present by mean ± SD. **P < 0.01, ***P < 0.001, ****P < 0.0001, and ns = no significance by One-way ANOVA in (c), (d), and (g). **P < 0.01 and ns = no significance by unpaired two sided student’s t-test in (e) and (j).

To determine the relevant producers of type 2 cytokines during chronic, latent infection, we then infected *Il4^KN^*^2^*^/+^* and *Il13^KN^*^4^*^/+^* mice to report endogenous IL-4 and IL-13 production, respectively^41,42^. Consistent with previous reports on cytokine-producing cells during *Nippostrongylus brasiliensis* infection, we observed that eosinophils and basophils exclusively express IL-4, ILC2s exclusively express IL-13, while T_H_2 cells produce both (Extended Data Fig. 3a and b)^42^. We next asked whether there was a dominant type 2 cytokine producing cell (as opposed to a redundant network) antagonizing fungal clearance. To address this, we infected *Epo^DTA^ (*eosinophil-deficient), *Mcpt8^Cre^ x Rosa26^LSL-DTA^ (*basophil-deficient), *Klrg1^Cre^ x Gata3^fl/fl^*(ILC2-deficient), *hCD2^Cre^ x Gata3^fl/fl^* (T_H_2-deficient), and paired controls with Δ*gcs1* to assess CFUs at day 35^43^. Eosinophil and basophil deficiency did not show a difference in pulmonary fungal burden, nor did ILC2 deficiency (Fig. 2e-g). On the other hand, T_H_2-deficient mice phenocopied the IL-4 and IL-13 double-knockout animals in displaying a dramatic reduction in pulmonary fungal burden (Fig. 2g), suggesting a dominant role for T_H_2 cells in establishing a type 2 inflammatory environment during latent infection that prevents fungal clearance.

To address the potential caveat that fungal infection fails to establish in the absence of type 2 inflammation, we asked whether temporal deletion of GATA3 during on ongoing infection would result in decreased pulmonary burdens. To test this, we infected *Rosa26^CreERT^*^2^ x *Gata3^fl/fl^* versus *Gata3^fl/fl^* control mice, started tamoxifen administration from 28 dpi, and performed CFU assays one week later at 35 dpi (Fig. 2h). Flow cytometry showed a successful ablation of GATA3 expression in T lymphocytes, which resulted in a significantly lower pulmonary fungal burden after just 7 days of deletion (Fig. 2i, j and Extended Data Fig. 3c), arguing that an ongoing type 2 response actively antagonizes fungal clearance.

### Spatial transcriptomics reveal the cellular architecture of cryptococcal granuloma in mouse models

Although our histological analysis showed structures consistent with granulomas, we wanted to obtain a more comprehensive understanding of the cellular and molecular microanatomy of these lesions. Therefore, we conducted spatial transcriptomics utilizing the 10x Genomics Xenium Prime 5K kit, which utilizes an *in situ* hybridization strategy to provide an accurate spatial gene expression pattern using a panel of 5006 mouse gene probes. This kit also contains multiple staining (DAPI, boundary (CD45, E-cadherin, and APT1A1), 18S, α-SMA, and vimentin) (Extended Data Fig. 4a) to aid in cell segmentation, thus enabling single-cell resolution for gene expression and allowing for annotation of cell types in addition to revealing spatial gene expression patterns. We harvested lung tissues from infected mice at 28, 35, 42, and 56 dpi and performed Xenium single-cell analysis to annotate cell types within the acquired sections. We were able to identify clusters pertaining to Mac/Mon, T lymphocytes, B lymphocytes, epithelial, endothelial, and stromal cells from all samples (Fig. 3a, b and Extended Data Fig. 4b). When we mapped these annotated cell types back to their spatial localization, we observed looser aggregates of immune cells at 28, 35, and 42 dpi, while the granulomatous structures were more organized at 56 dpi (Fig. 3b, bottom). Similar to mycobacterial granulomas in humans, we observed that murine cryptococcal granulomas had a ring-like myeloid core and a distinct T/B lymphocyte cuff (Fig. 3c), which is not usually found in murine Mtb infection^28^. Interestingly, epithelial cells were not observed in the granuloma core and were instead replaced by stromal cells that are likely heterogenous mixtures of fibroblasts (Fig. 3c). Consistent with highly organized microanatomy, we also observed spatially distinct chemokine expression (Fig. 3d). While there has been speculation that the lymphocyte cuff observed in Mtb granulomas is due to exclusion from the core, we found clear induction of *Cxcl13*, which guides B cells into secondary lymphoid follicles, in the region of the cuff, suggesting that lymphocytes may be actively recruited to the ring rather than excluded from the center^44^.

**Fig. 3:**
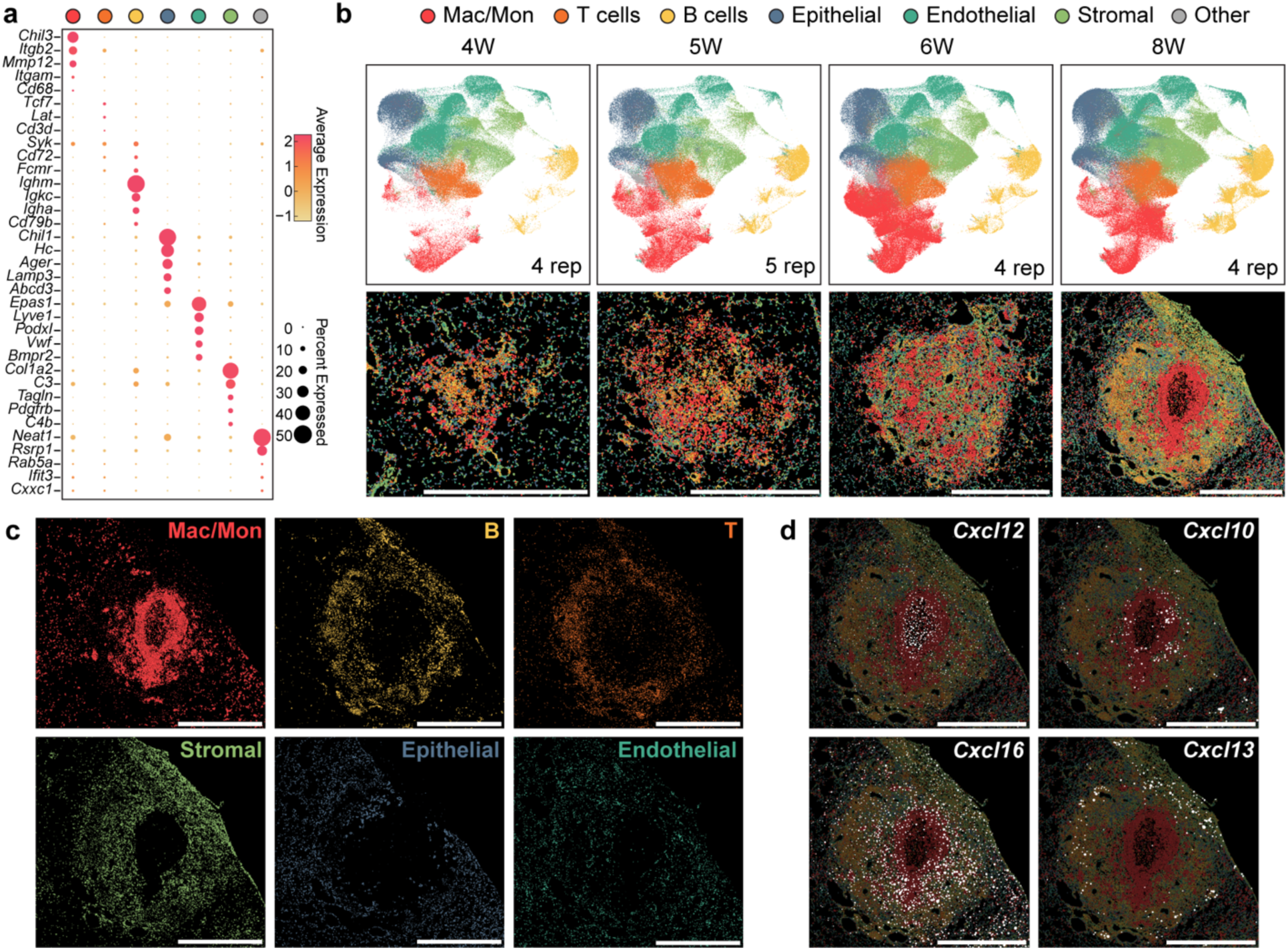
Spatial transcriptomics reveal the cellular architecture of cryptococcal granuloma. (a) Dot plots for top differentially expressed genes by different types of cells acquired by spatial transcriptomics. (b) Dynamics of different types of cells among time courses. Top, UMAP representation of merged images from all replicates at each time point. Bottom, representative images for spatial localization of cell types at each time point. (c) Images for different types of cells at 56 dpi. (d) Spatial expression of chemokines at 56 dpi. Scale bar, 1 mm.

### Type 2 signaling establishes an ARG1^+^ myeloid ring around the granuloma core

To further investigate the myeloid ring at the center of the granuloma, we were able to subset and re-cluster the Mac/Mon population into 4 clusters based on their featured gene expression (Fig. 4a, Extended Data Fig. 5a and b). Interestingly, each population within these 4 clusters of Mac/Mon displayed unique spatial localization (Fig. 4b). While the Mac/Mon4 cluster is largely excluded from the granuloma, Mac/Mon1, 2, and 3 were arranged in order from the interior to the outer layer of myeloid center. From analyses of cluster-defining genes, we found *Arg1* highly expressed in Mac/Mon3 (Fig. 3a). Spatially, *Arg1* expression covered the entire myeloid center forming a ring (Fig. 3c and d). Although ARG1 is well-known as a marker for AAM, it can also be induced under other conditions such as STAT3-activating cytokines or hypoxia^45–47^.

**Fig. 4:**
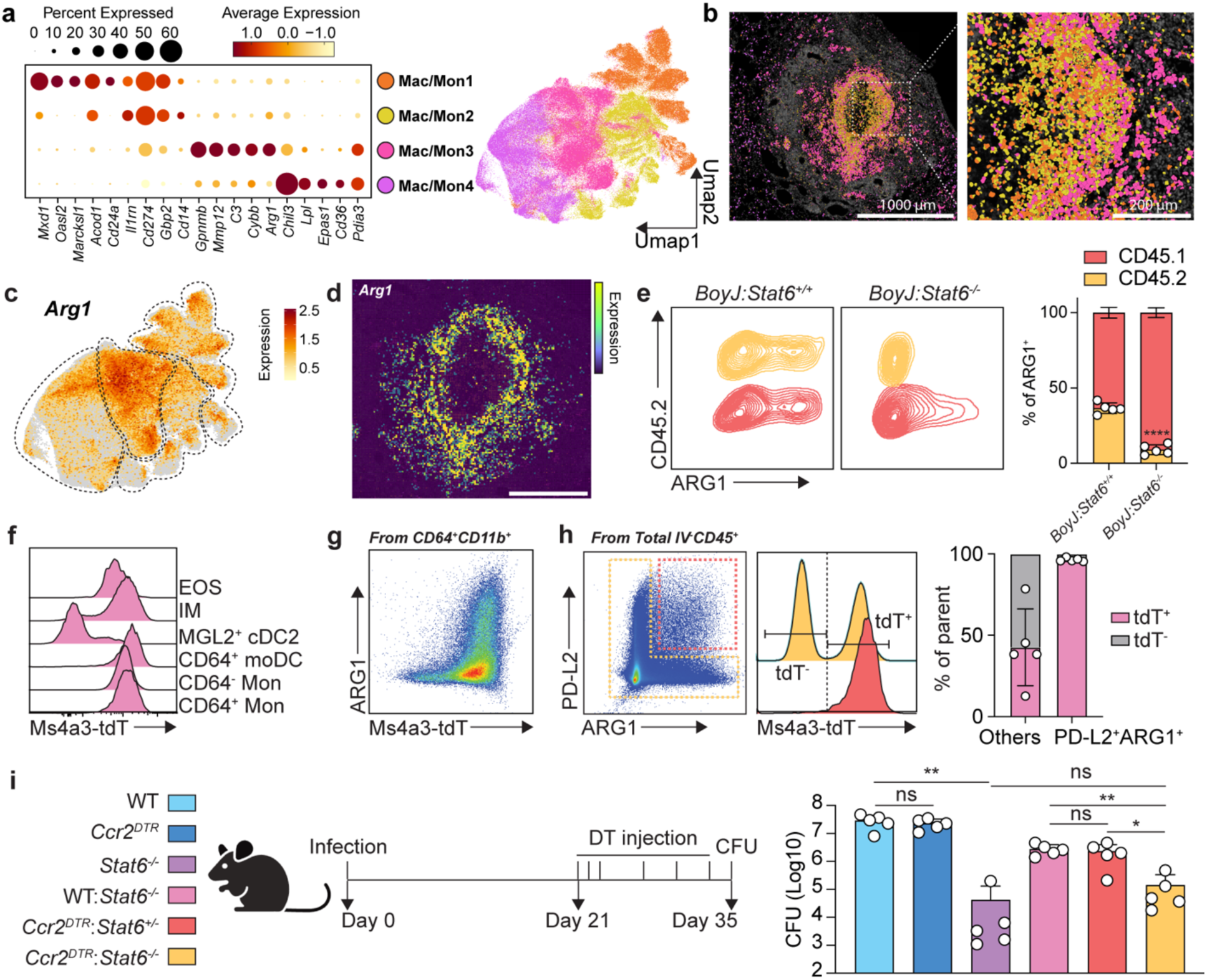
Type 2 signaling activates monocyte-derived myeloid cells to antagonize fungal clearance. (a-d) Macrophage/monocyte (Mac/Mon) subsets analysis. (a) Dot plot for top differentially expressed genes by 4 Mac/Mon clusters and UMAP representation of Mac/Mon clusters. (b) Spatial localization of 4 Mac/Mon clusters at 56 dpi. Scale bar, as indicated in figure. (c) Feature plot for *Arg1* expression in UMAP representation of Mac/Mon clusters. (d) Spatial *Arg1* expression at 56 dpi. Scale bar, 500 μm. (e) ARG1 expression in *BoyJ:Stat6^+/+^* or *BoyJ:Stat6^-/-^* mixed competitive chimeras. Left, flow plots; Right, statistics for the contribution of CD45.1 and CD45.2 bone marrows to ARG1 expressors. (f-h) Fate mapping by Ms4a3*^Cre^* x *Rosa26^LSL-tdTomato^*reporter mice at 35 dpi. (f) Histogram of tdTomato expression by myeloid cells. (g) Flow plot for ARG1 expression vs. tdTomato expression. (h) Analysis for tdTomato expression in ARG1+PD-L2+ cells and others. Left, flow plot for ARG1 vs. PD-L2; middle, histogram for tdT expression; right, statistics. (i) Stat6 specific depletion in CCR2+ cells. Left, experimental scheme; right, CFU analysis at 35 dpi. Data are present by mean ± SD. *P < 0.05, **P < 0.01, and ns = no significance by One-way ANOVA.

To determine whether ARG1 induction during *Cryptococcus* latent infection is due to cell-intrinsic STAT6 signaling, we generated mixed bone marrow chimeras whereby congenic markers can be utilized to recognize the genotype of cells competed in the same recipient. To minimize any potential decrease in fungal burden in *Stat6^-/-^*bone marrow mixes from cells that may gain increased killing capacity, we mixed 30% *Stat6^+/+^* (WT, CD45.2) or *Stat6^-/-^* (*Stat6* deficient, CD45.2) bone marrow with 70% WT bone marrow from *BoyJ* mice (WT, CD45.1). We then transplanted these mixes into irradiated recipients (WT, CD45.1/2) (Figure S5C). After reconstitution, we infected these chimeras and performed flow cytometry analysis. Within CD64^+^CD11b^+^ myeloid cells, which we found to represent the major ARG1 expressors, *Stat6^+/+^* cells expressed the same level of ARG1 compared to CD45.1 WT cells. In contrast, *Stat6^-/-^* cells lost almost all ARG1 expression, while CD45.1 WT cells in the same recipient still expressed ARG1, albeit to slightly diminished levels compared to the control mixes (Fig. 4e). We further gated on all ARG1^+^ myeloid cells to see the contribution from CD45.1 and CD45.2. The contribution of CD45.2 in *BoyJ:Stat6^+/+^*group was around 30%, whereas in *BoyJ:Stat6^-/-^* mixes the representation of knockout cells was significantly decreased. We further included PD-L2 as another type 2 marker^48^ and found the same intrinsic loss in *Stat6^-/-^* cells compared to WT cells (Extended Data Fig. 5d). These data suggest that the ARG1^+^ myeloid core is induced by type 2 cytokine signaling in a STAT6-dependent manner.

In summary, we identified that cryptococcal granulomas deriving from Δ*gcs1* latent infection are comprised by an inner myeloid core surrounded by ARG1+ type 2 responsive cells, an intermediate layer of stromal cells, and an outer cuff of T/B lymphocytes.

### Type 2 cytokines signal to monocyte-derived myeloid cells to antagonize fungal clearance

Next, we sought to investigate the ontogeny of the ARG1^+^ myeloid cells and whether these cells antagonize fungal clearance. Tissue myeloid cells can derive from embryonically seeded precursors that self-renew and persistent through adulthood, or from newly recruited bone marrow-derived monocytes that differentiate in the tissue^49,50^. To assess myeloid cell ontogeny during chronic *Cryptococcus* infection, we utilized *Ms4a3^Cre^* x *Rosa26^LSL-tdTomato^* fate mapper mice to see whether type 2 cytokine-responsive myeloid cells originate from adult bone marrow granulocyte monocyte progenitors (GMPs)^51^. During infection, the majority of ARG1 expressing cells were labeled by tdTomato (Fig. 4f and g). We further gated on ARG1^+^PD-L2^+^ cells from the entire resident immune cell population and compared the tdTomato expression with the remaining cells. We observed that almost all PD-L2^+^ARG1^+^ were labeled by tdTomato (Fig. 4h), arguing for an adult GMP monocyte-derived origin of these cells. To rule out contributions from classical dendritic cells (cDCs), since many of the CD11b^+^CD64^+^ cells in the lung are also CD11c^+^MHCII^+^, we infected *Zbtb46^Gfp^* reporter mice to see whether cDCs were contaminating our myeloid gates, especially the moDC gate^52^. Only ∼10% of CD64+ moDCs showed GFP expression, while as a positive control all classical DCs (cDC1 and cDC2) were GFP+ (Extended Data Fig. 5e and f). In summary, we identified that type 2 responsive myeloid cells during latent infection were mostly derived from GMPs with minimal contribution from cDCs.

We then asked whether STAT6 expression in monocyte-derived cells antagonized fungal clearance, presumably downstream of type 2 cytokine signals from T_H_2 cells. We mixed *Ccr2^DTR^* bone marrow with *Stat6^-/-^* bone marrow 1:1 and transplanted into irradiated recipients to specifically ablate STAT6 from the monocyte-derived compartment (Extended Data Fig. 5g). To control for losing 50% of monocytes, and for any dominant effect from 50% *Stat6^-/-^*bone marrow, we set up *Ccr2^DTR^*:*Stat6^+/-^* and WT:*Stat6^-/-^* control mixes. We started diphtheria toxin (DT) injection from 21 dpi to deplete type 2 signaling in CCR2^+^ cells after the initial T_H_2 peak (Fig. 4i). From this experiment, we found that *Stat6*-deletion in CCR2^+^ cells almost fully phenocopied *Stat6^-/-^* full chimeras in their reduction in lung fungal burden (Fig. 4i), suggesting that type 2 cytokines dominantly act through monocyte-derived myeloid cells to antagonize sterilizing immunity to *Cryptoccocus*.

### Type 2 immunity antagonizes extracellular killing of cryptococcus

In bacterial infection, the canonical model for type 1 immunity is that IFNγ signaling to macrophages increases their cell-autonomous killing capacity, thereby restricting intracellular pathogen replication^53^. For intracellular pathogens such as *Toxoplasma*, *Leishmania*, and *Salmonella*, published data argues that AAMs may function as an intracellular replication niche due to their dampened expression of antimicrobial effectors^54–56^. This argument is consistent with *in vitro* studies showing that STAT6 can repress STAT1 target genes^37^. Therefore, it is assumed that during fungal infection, type 2 signaling may also antagonize macrophage intracellular fungicidal capacity. To formally test whether this is the case during latent fungal infection, we again used a mixed chimera strategy to compare the intracellular fungal burden of WT and *Stat6*-deficient immune cells in the same microenvironment. We infected the *Stat6^+/+^*:BoyJ and *Stat6^-/-^*:BoyJ chimeras with Δ*gcs1*-mCherry reporter strain and gated out cryptococcus anti-capsule (glucuronoxylomannan, GXM) antibody positive yeast, which we presume to be extracellularly bound to host cells but not phagocytosed, as intracellular *Cryptococcus* should be shielded from the anti-GXM antibody. By gating on mCherry^+^GXM^-^ cells, we identified cells containing intracellular *Cryptococcus* (Fig. 5a). Within the immune compartment, the majority of these events were CD11b^+^CD64^+^ myeloid cells, which overlaps with the population we identified earlier as being type 2 cytokine responsive (Fig. 5b). If STAT6-deficient effector cells were more capable of killing intracellular *Cryptococcus*, the representation of CD45.2^+^ in *Cryptococcus*-containing effector cells should significantly decrease in the *BoyJ:Stat6^-/-^* chimeras but not in *BoyJ:Stat6^+/+^*. However, when we gated on total effector cells as well as *Cryptococcus*-containing effector cells and compared the CD45.2 representation within the two gates, there was no difference (Fig. 5c and d), suggesting identical intracellular killing capacity, or lack thereof, between WT and STAT6-deficient myeloid cells.

**Fig. 5:**
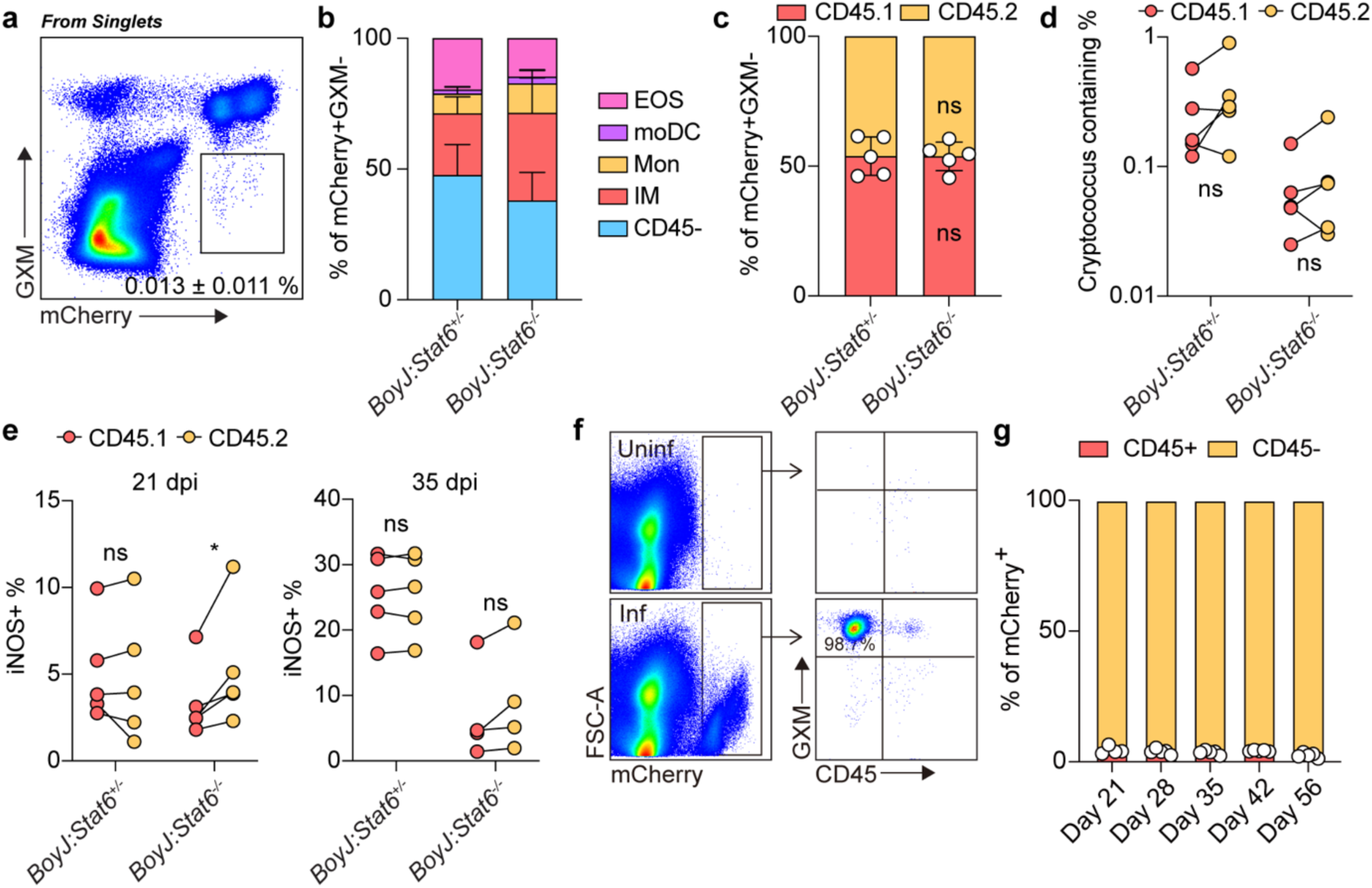
Type 2 signaling impairs extracellular fungal clearance. (a-d) *BoyJ:Stat6^+/-^* or *BoyJ:Stat6^-/-^*mixed competitive chimeras. (a) Flow plot for mCherry reporter and GXM staining. (b) Statistics for ratio of cells containing intracellular *Cryptococcus*. (c) The contribution of CD45.1 and CD45.2 to mCherry+GXM-CD64+CD11b+ myeloid cells. (d) The percent of *Cryptococcus* containing (mCherry+GXM-) cells in CD45.1 and CD45.2 CD64+CD11b+ myeloid cells. (e) The percent of iNOS+ cells in CD45.1 and CD45.2 CD64+CD11b+ myeloid cells. (f) Flow plots for mCherry+ cells and GXM vs. CD45 staining. (g) The percent of mCherry+ cells in CD45+ and CD45-cells. Data are present by mean ± SD. In (c), ns = no significance by unpaired student’s t-test. *P < 0.05 and ns = no significance by paired two sided student’s t-test in (d) and (e).

As STAT6 is reported to be able to repress induction of STAT1 target genes, we also analyzed whether iNOS production was affected by type 2 signaling. At both 21 dpi and 35 dpi in the mixed chimeras, we observed an unremarkable, albeit statistically significant at 21 dpi, increase in iNOS expression in *Stat6^-/^*^-^ myeloid cells (Fig. 5e). At 35 dpi, overall iNOS production by effector cells in *BoyJ:Stat6^-/-^* group was significantly lower than that in *BoyJ:Stat6^+/+^* group, likely due to the lower fungal burden caused by 30% *Stat6^-/-^* bone marrow mix (Extended Data Fig. 6a). These results suggest that *in vivo*, AAMs are not a replication niche for *Cryptococcus*, and the major mechanism whereby type 2 cytokines impair myeloid effector function is not repressing interferon-responsiveness.

Given that AAMs do not appear to be a *Cryptococcus* replication niche, we next asked whether the majority of the yeast were intracellular or extracellular. When assessing GXM staining on mCherry^+^ events (*Cryptococcus*), we noticed that most *Cryptococcus* yeasts during latent infection were not intracellular (Fig. 5a). Most yeasts were labeled by anti-GXM antibody and not associated with CD45^+^ cells (Fig. 5f and g). As a second complementary approach, we utilized live imaging on vibratome thick-section lung slices to observe fungal-immune interactions in real time. Consistent with the flow cytometry data, most *Cryptococcus* yeasts were outside of CD11b^+^, CD11c^+^, or CX3CR1^+^ myeloid cells (Video 1 and Extended Data Fig. 6f). We were occasionally able to observe overlap between CD11b and mCherry signals, which might represent intracellular *Cryptococcus* phagocytosed by CD11b+ cells. Still, these events were very rare (Video 2), consistent with the reported antiphagocytic role for the GXM capsule^57^. To make sure that the observations on the extracellular nature of *Cryptococcus* were not selective to the Δ*gcs1* mutant, we did the same STAT6-mixed chimera experiment infected with a WT KN99-mCherry strain, where we observed an identical phenotype (Extended Data Fig. 6b and c).

### STAT6 signaling drives myeloid cell exclusion away from *Cryptococcus*

As IL-4 has been previously shown to drive the proliferation of tissue-resident macrophages in the peritoneum and the pleura^58^, another possibility is that, rather than affecting the intrinsic polarization state of myeloid cells, type 2 cytokines may function to amplify the number of non-restrictive myeloid cells. However, in our mixed chimera experiments, we actually observed a slightly higher contribution of *Stat6*-deficient cells in the monocyte-derived myeloid cell compartment (Extended Data Fig. 6d and e), suggesting a potential repressive role of type 2 signaling in these cells.

From our spatial transcriptomic data, *Arg1*^+^ myeloid cells appeared to be localized further from the granuloma center than interferon-responsive cells. We wondered whether this spatial localization was driven by STAT6 signaling, as this might restrict their ability to mediate extracellular fungal clearance. To test this hypothesis, we generated *CD11c^Yfp^Stat6^-/-^*mice and then made 30%:70% mixed bone marrow chimeras (*CD11c^Yfp^*:WT vs *CD11c^Yfp^Stat6^-/-^*:WT) for imaging after infection (Fig. 6a). This system allowed us to see whether STAT6 intrinsically controlled myeloid cell positioning. Intriguingly, *CD11c^Yfp^Stat6^-/-^* cells were more localized within the granuloma core (Fig. 6b), suggesting that STAT6, at least partly, represses extracellular fungal killing by impairing the ability of myeloid cells to co-localize with infectious yeast.

**Fig. 6:**
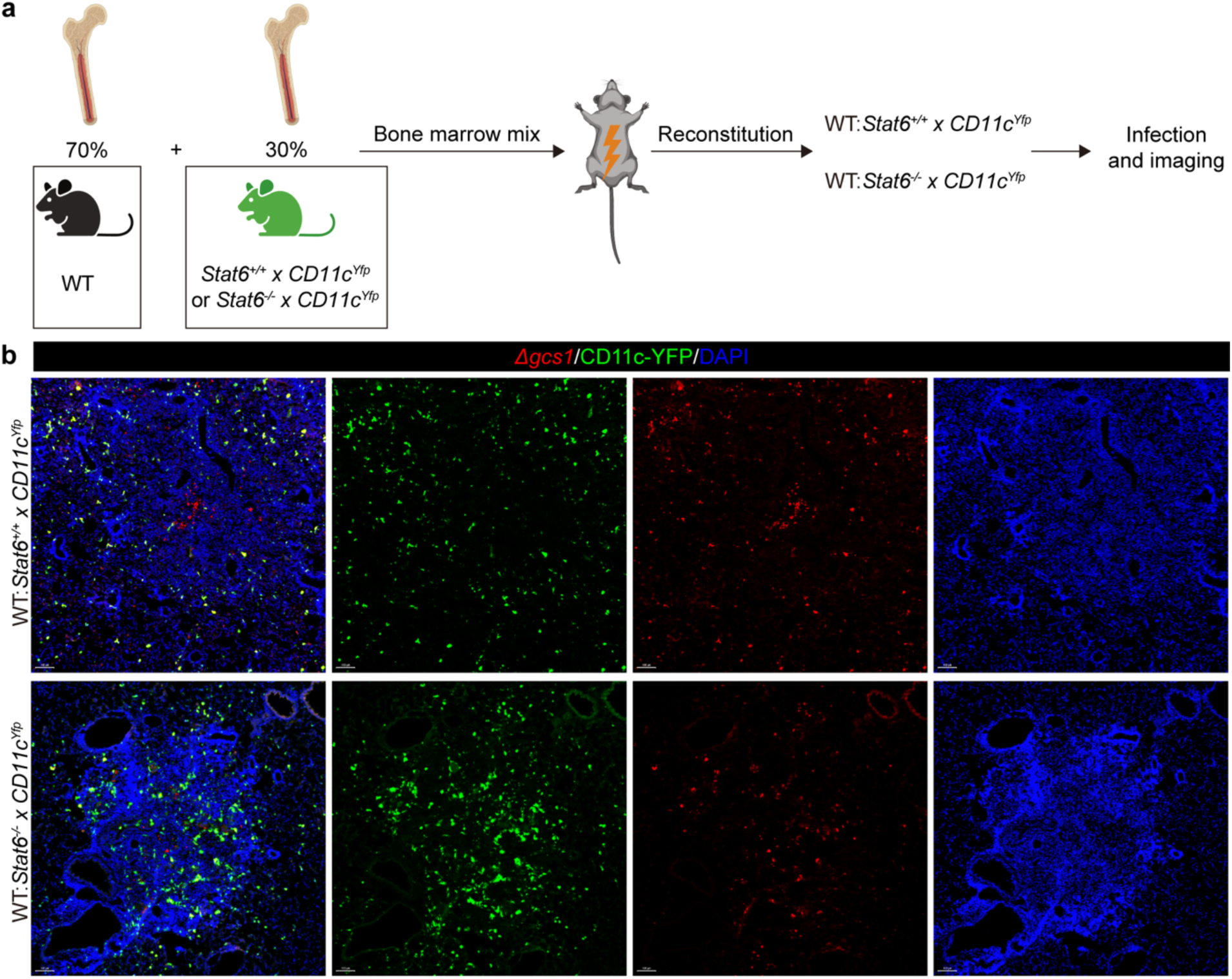
STAT6 signaling drives myeloid cell exclusion away from *Cryptococcus*. (a) Scheme and (b) representative images for STAT6 reporter mixed chimera (n=3). Scale bar, 100 μm.

## Discussion

In this study, we demonstrate that type 2 inflammation actively antagonizes sterilizing immunity to *Cryptoccocus* during non-lethal granulomatous infection. We show that the type 2 response is mainly established by T_H_2 cells via suppression of monocyte-derived myeloid cells in a manner that impairs extracellular fungal clearance. These observations are consistent with studies that have found detrimental roles for type 2 responses during acute lethal murine *Cryptococcus* infection, however the acute models do not appropriately mimic human infection^17,18^. While type 2 signaling deletion results in a lower pulmonary fungal burden during acute infection, it cannot significantly protect animals from mortality. Meanwhile, *Ifng^-/-^* and *Rag1^-/-^* mice have the same survival rate compared to WT mice (Extended Data Fig. 1a), suggesting the induction of a completely maladaptive T cell response during lethal infection, which is a very different infectious course compared to humans. Therefore, it is not a *priori* predicted that type 2 inflammation would play a role in scenarios where hosts do not succumb to chronic, subacute infection. Consistent with recent studies using clinical *Cryptococcus* isolates^9,59^, we find that the type 2 response is induced early but declines to a low level in this latent infection model. Our data argue that a long-lasting type 2 response can antagonize fungal clearance without causing fungal overgrowth that leads to host fatality. These findings raise the possibility of utilizing IL-4Rα blocking antibodies to pre-clear hosts of latent fungi before going on immunosuppressants for transplant surgery or chemotherapy.

During type 2 biased lethal serotype D *Cryptococcus* infection, it has been shown that *LysM^Cre^*x *Il4ra^fl/fl^*mice phenocopy *Il4ra* germline knockouts in displaying infection resistance^17^, arguing that macrophages are the major targets of type 2 cytokines that antagonize sterilizing immunity. However, *LysM^Cre^* has broad activity, being active in neutrophils, tissue resident macrophages, monocytes, some neurons, and airway epithelium^60–62^. Utilizing genetic fate mapping and genetic depletion, we demonstrate that monocyte-derived myeloid cells including IMs, moDCs, and monocytes are the major target cells of type 2 signaling that impair fungal clearance. Why AAMs, and type 2 immunity in general, are detrimental during fungal infection has been unclear. The most common argument is that since AAMs downregulate intracellular killing molecules (iNOS, etc), they may have impaired intrinsic fungicidal capacity, thus generating an intracellular replication niche. Utilizing a *Cryptococcus* reporter strain and extracellular capsule antibody staining, we were able to test this hypothesis *in vivo*. We did not observe a difference in intracellular fungicidal capacity between WT and *Stat6*-deficient macrophages. Instead, we find that most yeasts during both lethal and latent infection are extracellular, indicating that *Cryptococcus* clearance may happen in the extracellular space.

It has been reported that *Cryptococcus* can be rapidly internalized by alveolar macrophages (AMs) after intratracheal infection and extracellular predominance occurs by 24 hours post-infection^63^. In our study, we found that AMs did not significantly expand during latent infection (Extended Data Fig. 2b) and in our spatial transcriptomic analysis, we observed that cells in the Mac/Mon 4 cluster were mostly excluded by granuloma. These cells highly expressed *Chil3*, indicating an AM identity. Generally, we did not observe a significant contribution of the AM compartment to granulomas during latent infection and most immunosuppressive myeloid cells derived from monocytes. It has been shown that AMs have a higher phagocytic capacity than IMs^64^, which may explain the extracellular fungal dominance in granulomas. Our study opens new avenues in studying host clearance of *Cryptococcus* infection and raises the question of how IFNγ might provide protection against an extracellular pathogen. Our spatial transcriptomics data provide gene expression patterns at single-cell resolution within cryptococcal granulomas and reveal highly heterogenous myeloid cells in the granuloma center, some of which appear highly interferon-responsive based on expression of *Il1rn*, *Nos2* and *Acod1*. Deeper mechanistic dissection of these discrete Mac/Mon subsets may provide more insights into how myeloid cells directly or indirectly mediate extracellular fungicidal effects downstream of IFNγ. Our STAT6 mixed chimera data provides a potential mechanism that type 2 signaling may drive myeloid exclusion to impair their fungicidal capacity. Future studies should focus on how STAT6 signaling influences the localization of myeloid cells, and whether this migration is initiated or enhanced by IFNγ signaling.

Taken together, our results identify the cell types that produce and respond to type 2 cytokines during chronic granulomatous cryptococcal infection. We propose that in contrast to its previously described role in promoting intracellular fungal growth, type 2 cytokines impair the extracellular killing capacity of monocyte-derived cells. Additionally, type 2 cytokines do not appear to repress the capacity of myeloid cells to respond to IFNγ, albeit we cannot rule out that certain interferon-dependent effector functions are antagonized independent of transcriptional alterations. These findings open the door for the utilization of therapies targeting type 2 cytokine signaling as host-directed strategies for treating latent *Cryptococcus* infection.

## Data availability

All source data are available upon request.

## ACKNOWLEDGEMENTS

This work was funded by the Division of Intramural Research of NIAID/NIH (NIAID; ZIA-AI001364). We thank the NIAID animal facility staff, as well as O. Schwartz and S. Ganesan (NIAID Biological Imaging Facility). We thank Jeff Zhu for *hCD2^Cre^xGata3^fl/fl^*and *Klrg1^Cre^xGata3^fl/fl^* mice, Bill Petri for *Il13^-/-^*bone marrow, Yasmine Belkaid for *Ccr2^DTR^* mice, Roxane Tussiwand for *Zbtb46^GFP^* bone marrow. We thank Katrin Mayer-Barber, Niki Moutsopoulos, Hao Jin, Dan Barber, and Mihalis Lionakis for helpful discussions.

## AUTHOR CONTRIBUTIONS

Y.Z. and E.V.D. designed the study and experiments and wrote the original manuscript. Y.Z. performed the experiments, analyzed the data, and created the figures. M.J.P., C.U.O., and H.D. performed mouse survival and flow cytometry experiments. O.K. performed live microscopy imaging on lung slices. E.A. performed analysis of spatial transcriptomics data.

## FIGURES AND FIGURE LEGENDS

**Video 1: Live imaging of lung samples in *Cryptococcus* latent infection.**

**Video 2: Rare phagocytosis events in *Cryptococcus* latent infection by CD11b^+^ cells.**

**Extended Data Fig. 1:**
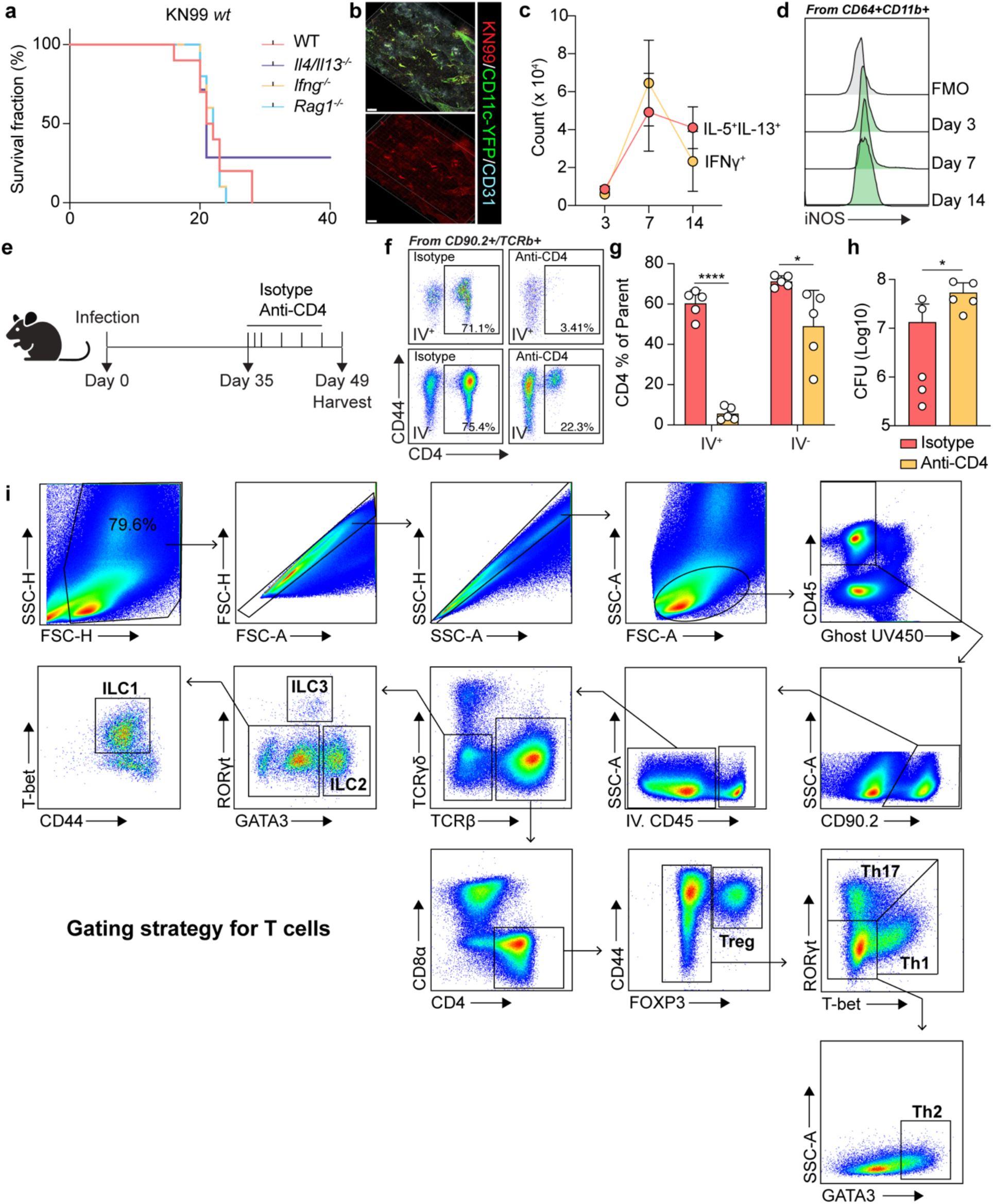
KN99 induces lethal infection and insufficient protective response in B6 mice. (a-d) KN99 induces a lethal infection without granuloma formation or effective iNOS induction. (a) Survival rate during KN99 infection (n=10). (b) Representative images for thick-tissue imaging of lungs from KN99 infected mice. Scale bar, 300 μm. (c) Statistics for cytokine production of restimulated CD4 T cells (n=5). (d) Histogram for iNOS expression in CD64+CD11b+ myeloid cells. (e-h) CD4 depletion in mice with KN99 *Δgcs1* infection. (e) Scheme for CD4 antibody treatment experiment. (f) Flow cytometry analysis of CD4 depletion in lung tissues at 49 dpi. (g) Statistics for efficiency of CD4 depletion. (h) CFU analysis. IV, intravenous labeling. (i) Gating strategy for T cell compartments. Data are present by mean ± SD.

**Extended Data Fig. 2:**
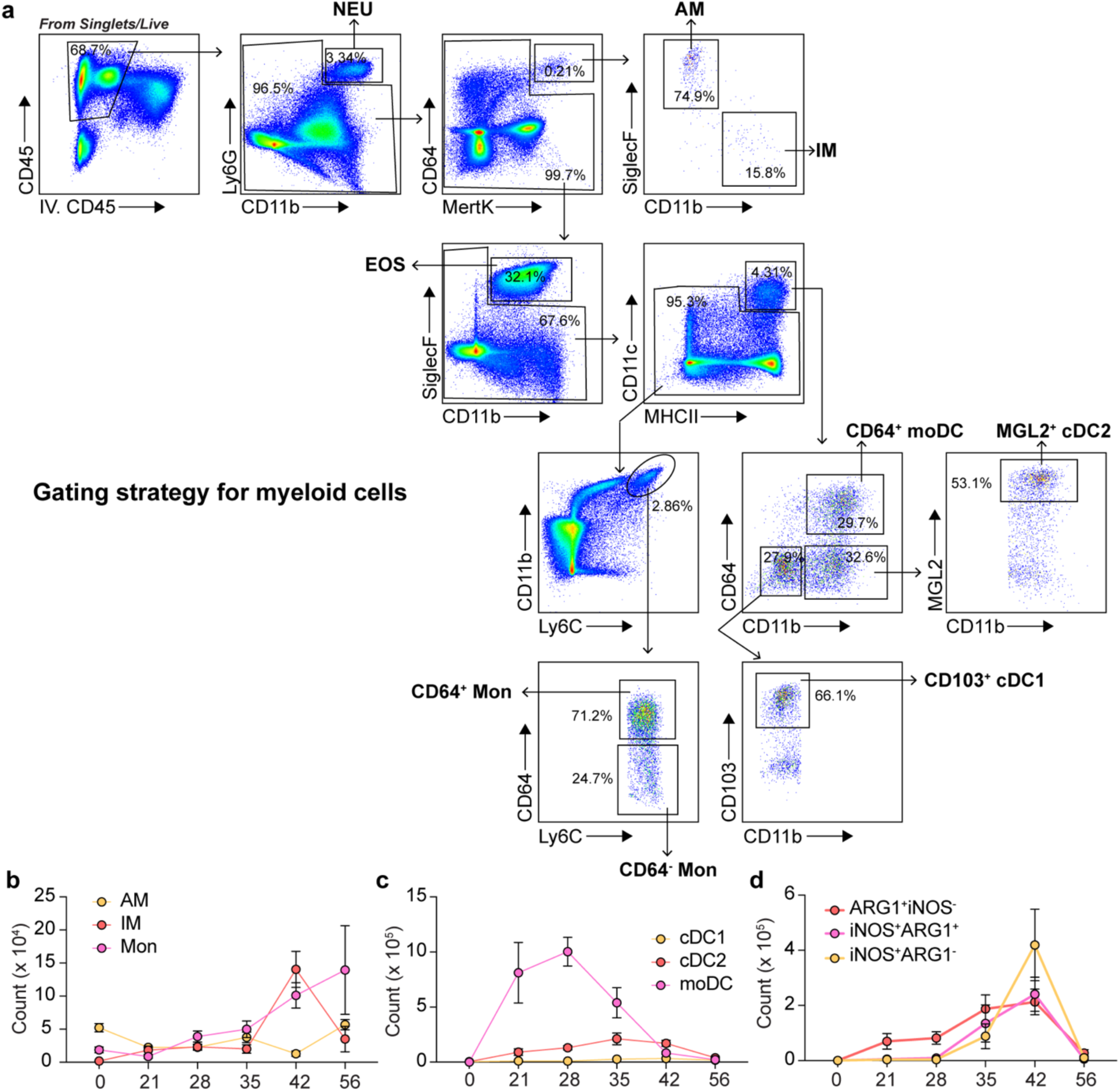
Myeloid cell expansion during KN99 *Δgcs1* infection. (a) Gating strategy for myeloid cells. NEU, neutrophil. AM, alveolar macrophage. IM, interstitial macrophage. EOS, eosinophil. Mon, monocyte. cDC, classical dendritic cell. moDC, monocyte-derived DC. (b-d) Statistics for myeloid cell expansion (n=5). Data are present by mean ± SD.

**Extended Data Fig. 3:**
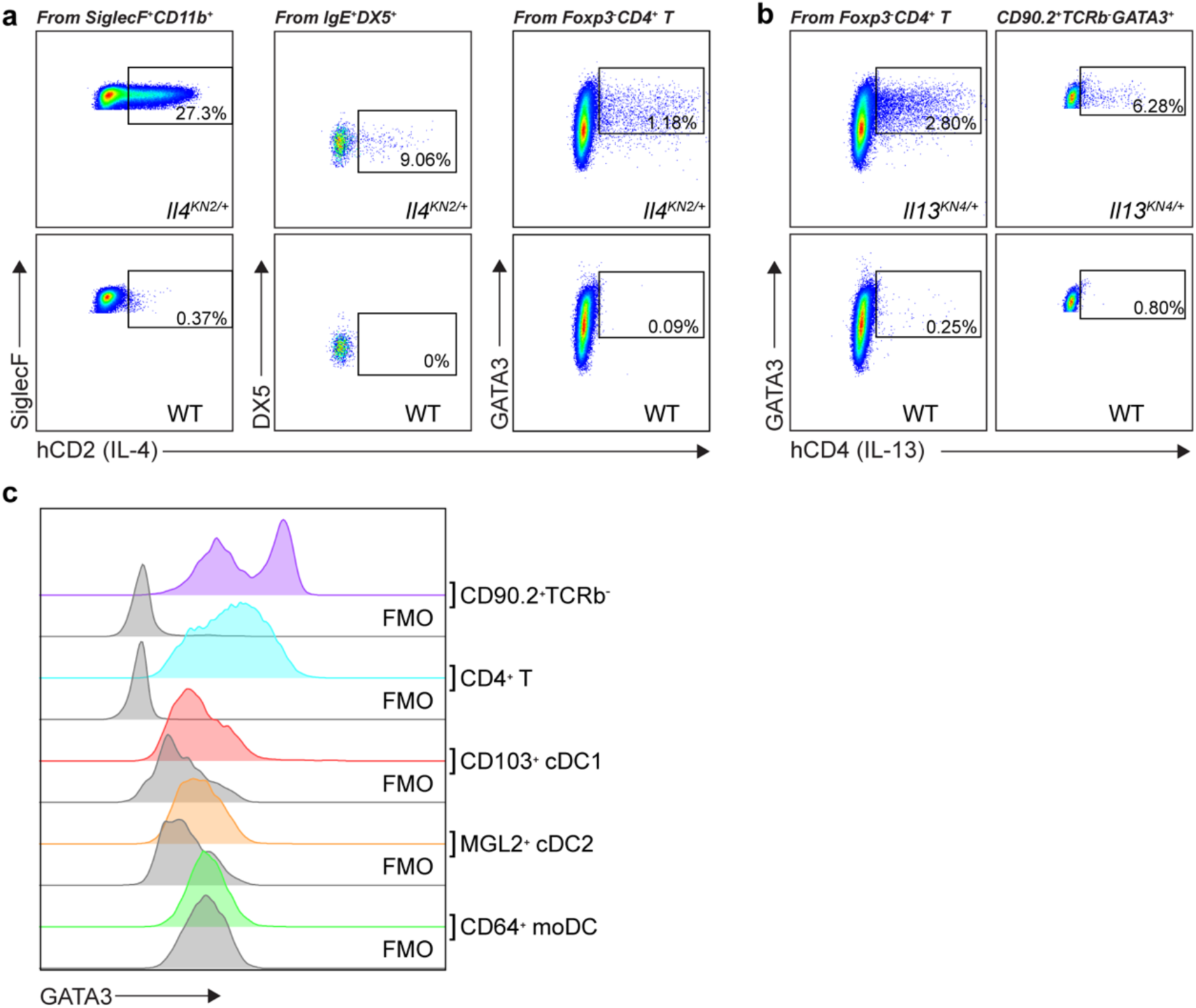
Type 2 cytokine producers during KN99 *Δgcs1* infection. (a) Flow plots for IL-4 production by eosinophils, basophils, and Th2 cells. (b) Flow plots for IL-13 production by Th2 cells and ILC2s. (c) GATA3 expression by T cells and DCs. FMO, fluorescence minus one control for GATA3 staining.

**Extended Data Fig. 4:**
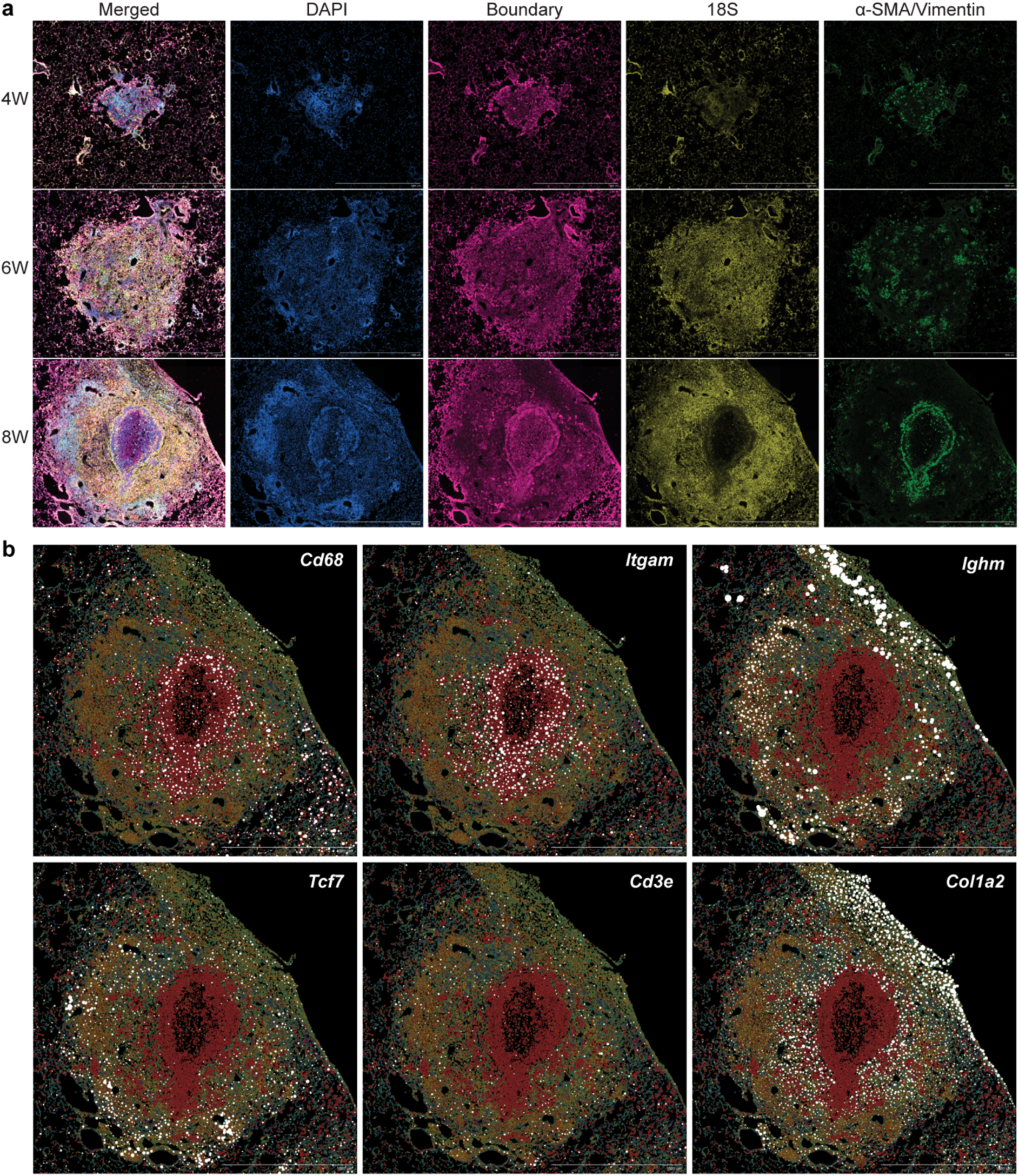
Spatial transcriptomics on cryptococcal granuloma. (a) Representative images for segmentation staining. (b) Lineage marker spatial expression at 56 dpi. Scale bar, 1 mm.

**Extended Data Fig. 5:**
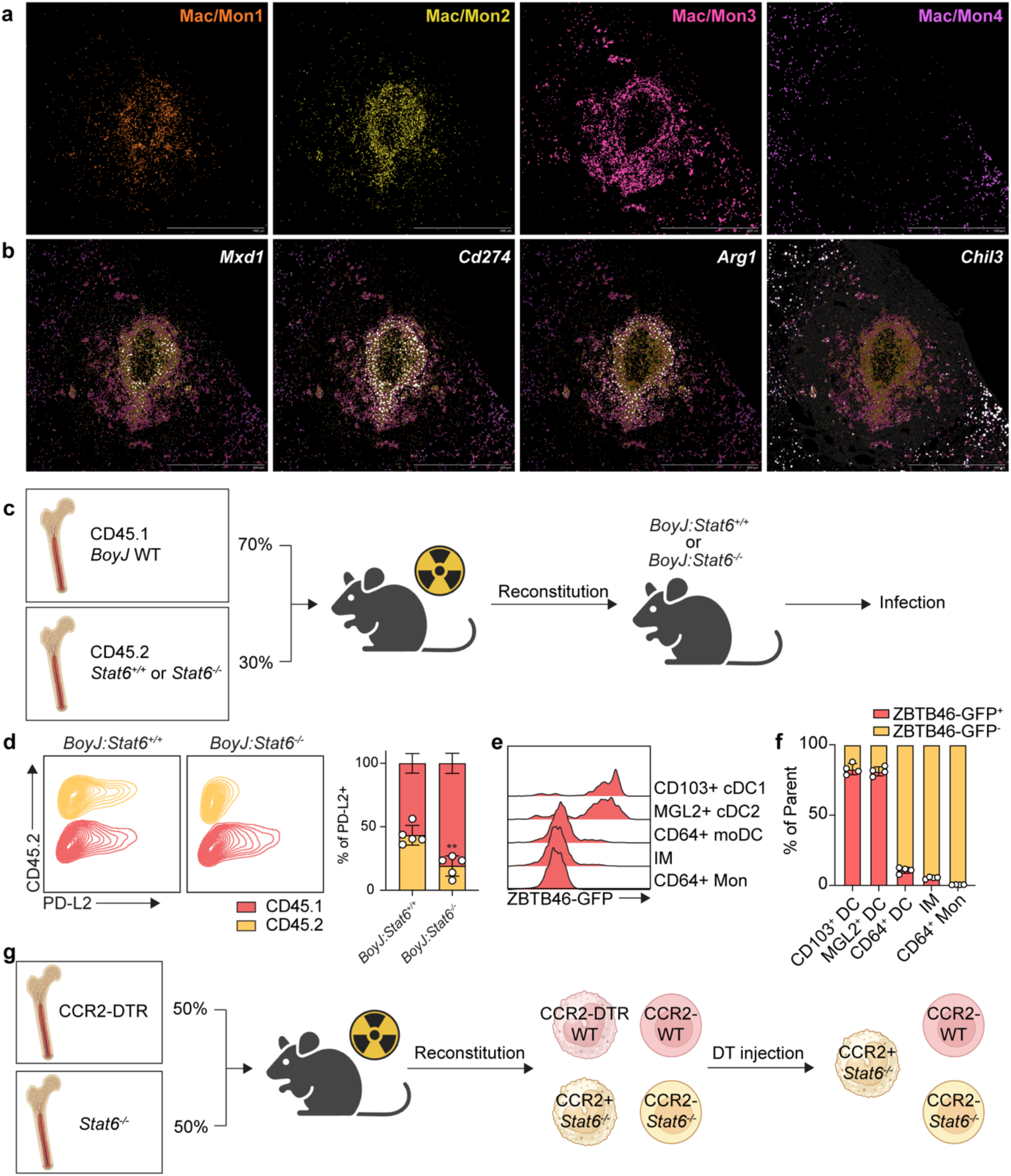
Subsets and origins of myeloid cells in granuloma center. (a) Spatial localization of 4 Mac/Mon clusters at 56 dpi. (b) Spatial expression of featured genes for 4 4 Mac/Mon clusters at 56 dpi. (c) Experimental scheme for mixed chimeras in Figure 4F. (d) PD-L2 expression in *BoyJ:Stat6^+/+^* or *BoyJ:Stat6^-/-^*mixed competitive chimeras. Left, flow plots; Right, statistics for the contribution of CD45.1 and CD45.2 bone marrow to PD-L2+ cells. (e, f) *Zbtb46^Gfp^* reporter expression at 35 dpi. (e) Histogram and (f) statistics of GFP expression by myeloid cells. (g) Experimental scheme for mixed bone marrow chimera in Figure 4J. Data are present by mean ± SD.

**Extended Data Fig. 6:**
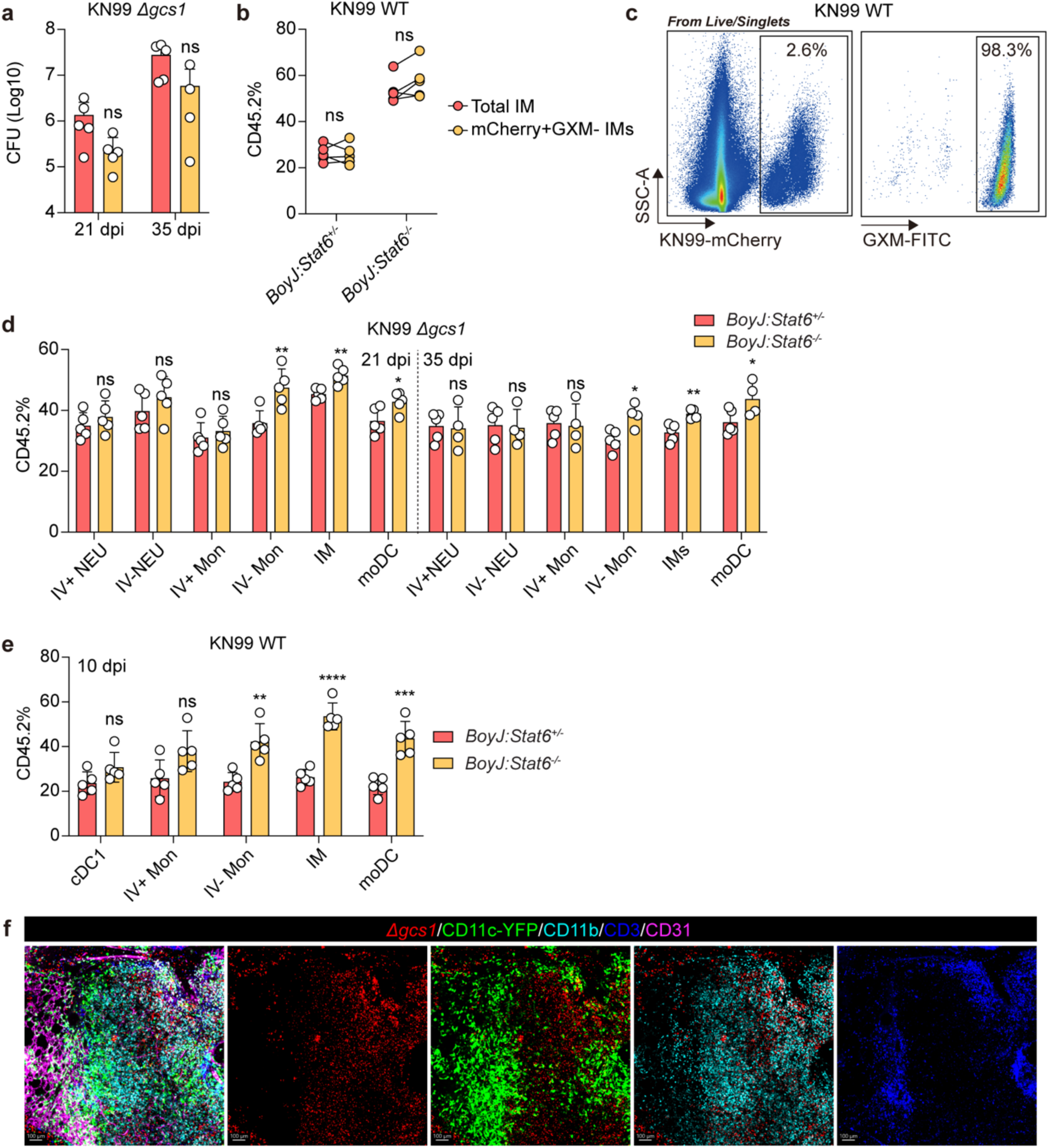
Type 2 signaling impairs extracellular fungal clearance. (a) CFU analysis in *BoyJ:Stat6^+/-^* or *BoyJ:Stat6^-/-^* mixed competitive chimeras during KN99 *Δgcs1* infection at 21 dpi and 35 dpi. (b) Statistics for the contribution of CD45.2 bone marrow derived cells to intracellular *Cryptococcus* containing IMs in KN99 infection. (c) Flow plots for mCherry reporter and GXM staining at 10 dpi in KN99 infection. |(d) CD45.2 ratio of different myeloid cells in *BoyJ:Stat6^+/-^* or *BoyJ:Stat6^-/-^* mixed competitive chimeras during KN99 *Δgcs1* infection at 21 dpi and 35 dpi. (e) CD45.2 ratio of different myeloid cells in *BoyJ:Stat6^+/-^* or *BoyJ:Stat6^-/-^* mixed competitive chimeras at 10 dpi during KN99 infection. (f) Representative images for live imaging of lung samples from infected CD11c-YFP mice at 70 dpi. Scale bar, 100 μm.

**Table S1:** information for antibodies.

| Antibodies | Source | Identifier |
| --- | --- | --- |
| AF488 anti-mouse ARG1 (clone A1EXF5) | eBioscience | Cat# 53-3697-80 |
| AF488 anti-mouse IgE (clone RME-1) | BioLegend | Cat# 406909 |
| AF488 anti-GFP (polyclonal) | ThermoFisher | Cat# A-21311 |
| AF647 anti-mouse Ly6G (clone 1A8) | BioLegend | Cat# 127609 |
| AF700 anti-mouse CD31 (clone 30090) | BioLegend | Cat# 102444 |
| AF700 anti-mouse EpCAM (clone G8.8) | BioLegend | Cat# 118240 |
| APC anti-mouse CD45 (clone 30-F11) | BioLegend | Cat# 103112 |
| APC-Cy7 anti-mouse CD11c (clone N418) | BioLegend | Cat# 117324 |
| APC-Cy7 anti-mouse CD49b (clone DX5) | BioLegend | Cat# 108919 |
| APC-Cy7 anti-mouse TCRβ (clone H67-597) | BioLegend | Cat# 109219 |
| APC-Cy anti-mouse IL-17A (clone TC11-18H10.1) | BioLegend | Cat# 506940 |
| BB700 anti-mouse GATA3 (clone L50823) | BD Bioscience | Cat# 566643 |
| BV421 anti-mouse CD3 (clone 17A2) | BioLegend | Cat# 100227 |
| BV421 anti-mouse IL-5 (clone TRFK5) | BioLegend | Cat# 504311 |
| BV421 anti-mouse Foxp3 (clone MF-14) | BioLegend | Cat# 126419 |
| BV421 anti-mouse PD-L2 (clone TY25) | BioLegend | Cat# 107219 |
| BV510 anti-mouse CD103 (clone 2E7) | BioLegend | Cat# 121423 |
| BV510 anti-mouse CD45.1 (clone A20) | BD Bioscience | Cat# 565278 |
| BV510 anti-mouse CD90.2 (clone 30-H12) | BioLegend | Cat# 105335 |
| BV605 anti-mouse CD4 (clone GK1.5) | BioLegend | Cat# 100451 |
| BV605 anti-mouse CD4 (clone RM4-4) | BioLegend | Cat# 116027 |
| BV605 anti-mouse MertK (clone 2B10C42) | BioLegend | Cat# 151517 |
| BV650 anti-mouse Ki67 (clone 11F6) | BioLegend | Cat# 151215 |
| BV711 anti-mouse CD69 (clone H1.2F3) | BioLegend | Cat# 104537 |
| BV711 anti-mouse Ly6C (clone HK1.4) | BioLegend | Cat# 128037 |
| BV711 anti-mouse IFNγ (clone XMG1.2) | BD Bioscience | Cat# 563736 |
| BV785 anti-mouse KLRG1 (clone 2F1/KLRG1) | BioLegend | Cat# 138429 |
| BV786 anti-mouse CD44 (clone IM7) | BD Bioscience | Cat# 563736 |
| BV786 anti-mouse SiglecF (clone E50-2440) | BD Bioscience | Cat# 740956 |
| FITC anti-mouse CD8 (clone S18018E) | BioLegend | Cat# 162313 |
| FITC anti-mouse IgG1 (clone RMG1-1) | BioLegend | Cat# 406605 |
| PE anti-human CD2 (clone RPA-2.10) | BioLegend | Cat# 300207 |
| PE anti-human CD4 (clone RPA-T4) | BioLegend | Cat# 300508 |
| PE anti-mouse T-bet (clone 4B10) | BioLegend | Cat# 644809 |
| PE-Cy5 anti-mouse CD64 (clone X54-5/7.1) | BioLegend | Cat# 139332 |
| PE-Cy7 anti-mouse CD11c (clone N418) | BioLegend | Cat# 117317 |
| PE-Cy7 anti-mouse iNOS (clone CXNFT) | eBioscience | Cat# 25-5920-80 |
| PE-Cy7 anti-mouse Ly6G (clone 1A8) | BioLegend | Cat# 127617 |
| PE-Cy7 anti-mouse CD301b (MGL2) (clone URA-1) | BioLegend | Cat# 146807 |
| PE-Cy7 anti-mouse RORγt (clone B2D) | eBioscience | Cat# 25-6981-82 |
| PerCP-Cy5.5 anti-mouse MHCII (clone M5/114.15.2) | BioLegend | Cat# 107625 |
| R718 anti-mouse IL-13 (clone W19-895) | BD Bioscience | Cat# 569945 |
| Anti-Glucuronoxylomannan (GXM) (clone 18B7) | Sigma-Aldrich | Cat# MABF2069 |
| InVivoMab anti-mouse CD4 (clone GK1.5) | Bio X Cell | Cat# BE0003-1 |
| InVivoMab rat IgG2b isotype control (clone LTF-2) | Bio X Cell | Cat# BE0090 |

## Method

### Animal experiments

Animal experiments were carried out at 14DNR in compliance with NIAID’s Animal Care and Use Committee’s guidance.

### *Cryptococcus* intranasal infection

Cryptococcus was obtained from frozen glycerol stocks and then cultured on a yeast extract peptone-dextrose (YPD) plate for two days, followed by overnight culture at 30 °C in YPD broth with shaking. Mice were anesthetized by 3% isoflurane and infected intranasally with 5×10^4^ CFU yeast in 25uL 1X PBS. Mice were monitored until recovery from anesthesia and were placed back to cages.

### Histological analysis

Lung samples were collected and fixed in 4 % PFA. Sectioning and H&E staining were performed and evaluated by veterinary pathologists at Infectious Disease Pathogenesis Section at NIAID. For immunofluorescent staining, euthanized and perfused mice were inflated with OCT via the trachea, and lung were collected and fast frozen. Frozen sections were made using a CryoStat into 8 μm thickness and air dried for 1 hour. Sections were then fixed with 4% PFA again and blocked with 5% BSA followed by hydrogen peroxide treatment. Fluorophore-conjugated antibodies were incubated at 4 °C overnight. Sections were mounted on slides for the imaging by Leica SP8 conjugated with 690 nm laser at the Bioimaging Research Technologies Branch at NIAID.

### Thick-tissue imaging

Mice were euthanized and perfused with 20 ml 1X PBS followed by 20 ml 4% PFA. Inflation with 2 % low-melting agarose was then performed for thick tissue sectioning using a Vibrating Blade Microtome. 500 μm sections were made and processed with Ce3D Tissue Clearing buffer set according to commercial instructions. Briefly, sections were permeabilized at room temperature for 2 days with gentle shaking followed by antibody staining for another 2 days with gentle shaking. Sections were washed 3 times by washing buffer during next 24 hours and then cleared by clearing solution for overnight. Cleared sections were mounted with clearing solution in 4 spacers (120 μm thickness for each) on slides for imaging by Leica SP8 conjugated with 690 nm laser at Bioimaging Research Technologies Branch at NIAID.

### Colony forming unit assay

Lung samples were collected and homogenized in sterilized water followed by serial dilution. KN99 *Δgcs1* contains a nourseothricin-resistance cassette. Therefore, all samples for CFU were plated on nourseothricin-containing YPD plates for fungal burden detection.

### CD4 antibody treatment

Anti-mouse CD4 antibody (GK1.5, BioXCell) or its isotype, rat anti-KLH IgG2b, were injected intraperitoneally with an initial 2 doses of 100 μg per mouse and rest being 4 doses of 200 μg per mouse.

### Flow cytometry

All mice for flow cytometry analysis in this study received retroorbital injection of 3 μg CD45 antibody 3 minutes before euthanasia to label intravascular cells. For lung samples, tissues were collected directly into 5 ml digestion buffer (HBSS with 1 mg/ml collagenase II, 1 mg/ml dispase, and 10 μg/ml DNAse I) and then digested using a GentleMACS. Digestion was stopped by adding 5 ml flow buffer (PBS with 2% FBS and 2 mM EDTA) on ice. Samples were immediately filtered using 70um strainers and centrifuged to remove digestion buffer. Red blood cells were removed by ACK lysis buffer for 5 min on ice and cells were then spun, washed, and aliquoted in 96-well round bottom plates for staining. Live/dead staining was performed for 10 min in PBS containing Fc Block before protein staining. Surface proteins were stained for 20 min on ice and then cells were acquired using a Cytek Aurora spectral flow cytometer. Intracellular and intranuclear proteins were stained with Transcription Factor Staining Buffer set. Briefly, cells were fixed and permeabilized on ice for 1 hour and stained in permeabilization buffer containing antibodies for 1 hour at room temperature. Cells were washed and resuspended in flow buffer for acquisition. T cell restimulation was performed for cytokine analysis. Collected cells were cultured in RPMI 1640 culture medium with PMA (50 ng/ml) and ionomycin (500 ng/ml) for 4 hours and then processed for staining. For GXM staining, cells were stained with other antibodies together with anti-GXM antibody (isotype: mouse IgG1) for surface staining for 20 min on ice. Then, after three times washing, cells were stained with FITC anti-mouse IgG1. Data were analyzed using FlowJo.

### Survival rate

Survival experiment was conducted by the coordination with animal facility 14DNR at NIAID and was monitored by veterinarians at Comparative Medicine Branch, NIAID. Endpoint was set up when the loss of body weight reached 20% compared to starting point. Animals were euthanized at the endpoint.

### Bone marrow chimera

Mice were irradiated at 700 rads in two doses spaced 3hrs apart by 14DNR at NIAID. Bone marrow was prepared and injected the same day at least one hour after the final irradiation dose. Mice were monitored during 6-week reconstitution. Bleeding was performed to determine reconstitution efficiency.

### GATA3 temporal depletion

Tamoxifen was administered by oral gavage with the dose of 75 mg/kg.

### Spatial transcriptomics

Euthanized mice were perfused with 20 ml PBS followed by 20 ml 4% PFA and then inflated with OCT from trachea. Lung tissues were collected and immediately fast frozen for storage at -80 °C. Sections were made with CrypStat by 10 μm directly on slides provided by 10X Genomics. Xenium Prime 5K kit was utilized for spatial transcriptomics according to commercial instructions. Briefly, sections were hybridized with priming oligos and then washed by RNase to release the RNA strand, followed by polishing to allow probe hybridization for the target RNA. We did not add customized probes on pre-designed mouse pan tissue & pathways panel containing 5006 probes. Unbound probes were washed away, and ligation was performed to ensure probe specificity by generating a circular DNA from hybridized probe, which was enzymatically amplified subsequently. Cell segmentation staining and autofluorescence quenching were performed before loading slides into Xenium Analyzer at NIAMS.

The output from the Xenium analyzer was processed with Seurat version 5.1.0.9006^65^. First, the Xenium data folder from each image was loaded onto Seurat with the ’LoadXenium’ function and the segmentations parameter set to ’cell’. Cells without feature counts (nCount_Xenium > 0) were subsequently filtered from the dataset. Each image contained a single section except for one, whose Seurat object was then split into two separate objects based on tissue coordinates using the ’Crop’ and ’subset’ functions. Next, all images were combined into a single Seurat object using ’merge’, and low-quality cells containing less than 3 Xenium features and 6 counts (nFeature_Xenium < 3 & nCount_Xenium < 6). Finally, data was normalized using the SCTransform function^66^.

For analysis, the dimensionality of the dataset was determined by running the ’RunPCA’ and ’ElbowPlot’ functions, and 25 dimensions were used in subsequent steps. Dimensionality reduction and clustering was performed with ’RunUMAP’, ’FindNeighbors’, and ’FindClusters’, with a range of resolutions. A clustering based on a resolution of 0.3 was chosen to capture the biological diversity in the dataset, resulting in 19 clusters. To perform cluster annotation, the differentially expressed genes per cluster were calculated using the ’PrepSCTFindMarker’ and ’FindAllMarkers’ (min.pct = 0.01) functions. Based on these genes and on visualization of cell type specific markers these 19 clusters were annotated into 7 major cell types, which were subsequently used for visualization on the Xenium Explorer (10X Genomics). Finally, these major cell types and the genes used in cluster annotation were visualized using Seurat.

For further analysis of macrophage/monocyte populations from the Xenium dataset, cells annotated as macrophage/monocyte were subset into a new Seurat object. Data processing an analysis was performed as described above for the entire dataset (starting at the normalization step with SCTransform) with the following parameters: 30 dimensions and resolution of 0.2, resulting in 22 macrophage/monocyte clusters. Subsequently these 22 clusters were grouped into four macrophage/monocyte subsets based on shared gene expression patterns and temporal occurrence.

### Diphtheria toxin (DT) treatment

DT was injected intraperitoneally with the dose of 100 ng per mouse.

### Live lung tissue imaging

Mice were euthanized using 5% Isoflurane at induction chamber. After euthanasia, mouse lungs were inflated with 1.5 % of Sea Kem agarose in phenol-red free RPMI at 37 °C. Inflated tissues were kept on ice, in 1 % FBS in PBS, and sliced into 300-350 µm sections using Leica VT1200 S Vibrating Blade Microtome (Leica Microsystems), in ice-cold PBS. Tissue sections were stained with fluorescently labeled antibody cocktail of choice for 2-6 h in the 37 °C incubator. After staining sections were washed 3 times and cultured in complete lymphocyte medium (Phenol Red-free RPMI supplemented with 20 % FBS, 25 mM HEPES, 50 μM β-ME, 1 % Pen/Strep/L-Glu and 1 % Sodium Pyruvate) in humidified incubator at 37°C. Tissues were allowed to completely recover for 12 h prior to recording time-lapse video. Sections were held down with tissue anchors (Warner Instruments) in 2-well imaging chambers and imaged using Leica DIVE inverted 5 channel confocal microscope equipped with an Environmental Chamber (NIH Division of Scientific Equipment and Instrumentation Services) to maintain 37 °C and 5 % CO_2_. Microscope configuration was set up for four-dimensional analysis (x,y,z,t) of cell segregation and migration within tissue sections. Diode laser for 405 nm excitation; Argon laser for 488 and 514 nm excitation, DPSS laser for 561; and HeNe lasers for 594 and 633 nm excitation wavelengths were tuned to minimal power (between 0.1 and 2 %). Z stacks of images were collected (10 – 50 µm). Mosaic images of lung sections were generated by acquiring multiple Z stacks using motorized stage to cover the whole section area and assembled into a tiled image using LAS X (Leica Microsystems) software. For time-lapse analysis of cell migration, tiled Z-stacks were collected over time (1 to 4 h). Post-acquisition mages were processed using Imaris software.

---

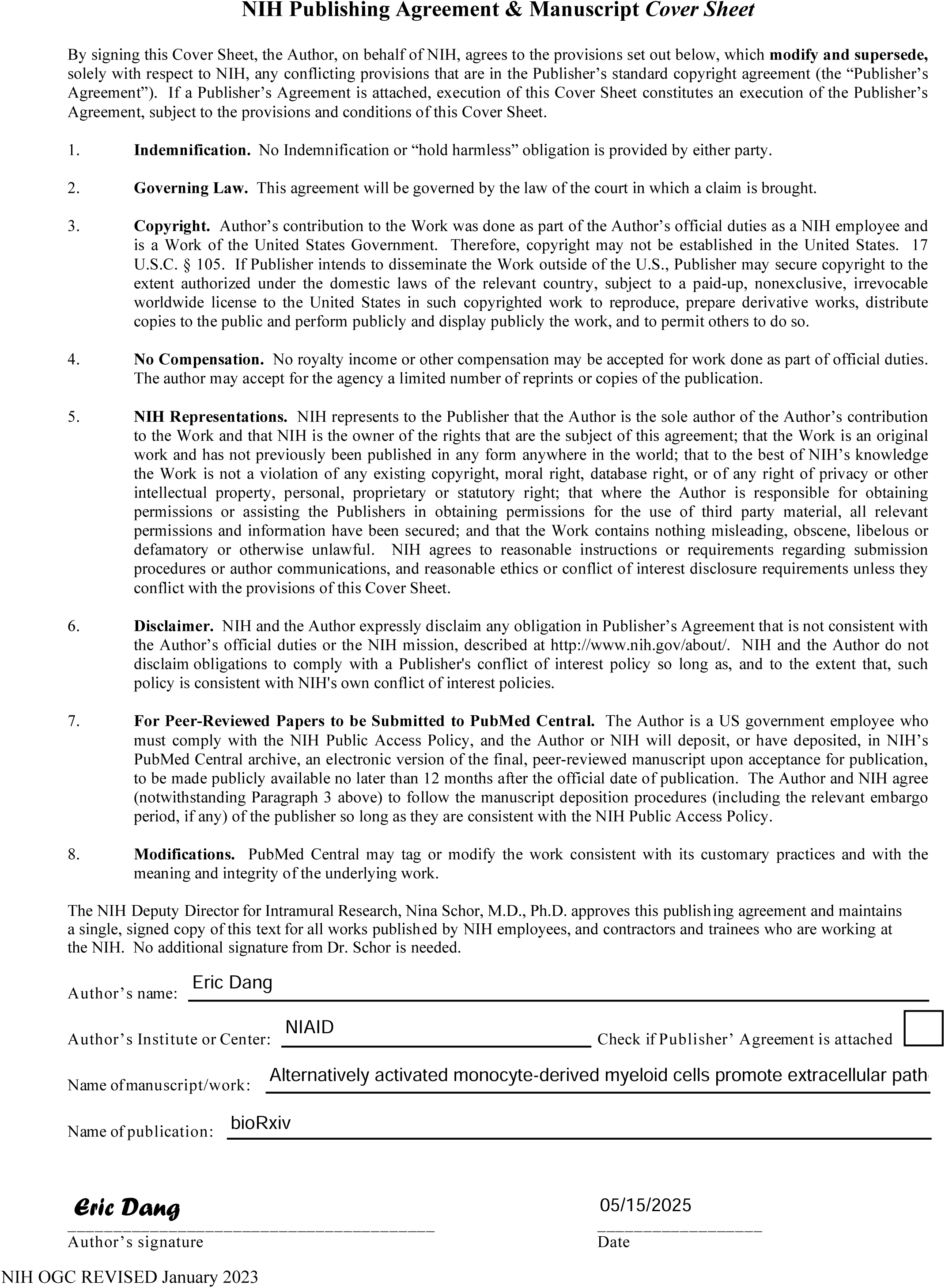

